# Interplay between gene nucleotide composition bias and splicing

**DOI:** 10.1101/605832

**Authors:** Sébastien Lemaire, Nicolas Fontrodona, Jean-Baptiste Claude, Hélène Polvèche, Fabien Aubé, Laurent Modolo, Cyril F. Bourgeois, Franck Mortreux, Didier Auboeuf

**Affiliations:** Univ Lyon, ENS de Lyon, Univ Claude Bernard, CNRS UMR 5239, INSERM U1210, Laboratory of Biology and Modelling of the Cell, 46 Allée d’Italie Site Jacques Monod, F-69007, Lyon, France; CECS, I-Stem, F-91100, Corbeil-Essonnes, France; LBMC biocomputing center, CNRS UMR 5239, INSERM U1210, 46 Allée d’Italie Site Jacques Monod, F-69007, Lyon, France

**Keywords:** splicing, genomic, chromatin organization, nucleotide composition bias

## Abstract

To characterize the rules governing exon recognition during splicing, we analysed dozens of RNA-seq datasets and identified ~3,200 GC-rich exons and ~4,000 AT-rich exons whose inclusion depends on different sets of splicing factors. We show that GC-rich exons have predicted RNA secondary structures at 5’-ss, and are dependent on U1 snRNP–associated proteins. In contrast, AT-rich exons have a large number of branchpoints and SF1-or U2AF2-binding sites and are dependent on U2 snRNP–associated proteins. Nucleotide composition bias also influences local chromatin organization, with consequences for exon recognition during splicing. As the GC content of exons correlates with that of their hosting genes, isochores and topologically-associated domains, we propose that regional nucleotide composition bias leaves a local footprint at the exon level and induces constraints during splicing that can be alleviated by local chromatin organization and recruitment of specific splicing factors. Therefore, nucleotide composition bias establishes a direct link between genome organization and local regulatory processes, like alternative splicing.

## Introduction

Most eukaryotic genes comprise both exons and introns. Introns are defined at their 5’-end by the 5’ splicing site (ss), which interacts with the U1 snRNA, and at their 3’-end, by the branchpoint (BP; recognized by SF1), the polypyrimidine (Py) tract (recognized by U2AF2 or U2AF65) and the 3’ ss (recognized by U2AF1 or U2AF35)^1^. SF1 and U2AF2 allow the recruitment of the U2 snRNP, which contains the U2 snRNA that interacts with the BP^1^. In addition to linear sequences (e.g., the Py tract), the secondary structures of RNA play important roles in splicing. For example, secondary structures at the 5’ ss can hinder the interactions between the 5’ ss and U1 snRNA^2,3^, and secondary structures at the 3’-end of short introns can replace the need for U2AF2^4,5^. Splicing signals are short, degenerate sequences, and exons are much smaller than introns. How then are exons precisely defined? How are *bona fide* splicing signals distinguished from pseudo-signals or decoy signals? These questions have been intensively researched but still remain open.

Many (if not all) exons require a variety of splicing factors to be defined. Splicing factors that belong to different families of RNA binding proteins, such as the SR and hnRNP families, bind to short degenerate motifs either in exons or introns of pre-mRNAs^6^. Splicing factor binding sites are low-complexity sequences comprising either the same nucleotide or dinucleotide^7–9^. Splicing factors modulate the recruitment of different spliceosome-associated components^6,10^.

Spliceosome assembly and the splicing process occurs mostly during transcription^10,11^. In this setting, the velocity of RNA polymerase II (RNAPII) influences exon recognition in a complex manner, as speeding up transcription elongation can either enhance or repress exon inclusion^12^. RNAPII velocity is in turn influenced by the local chromatin organization, such as the presence of nucleosomes^10,11^. Nucleosomes are preferentially positioned on exons because exons have a higher GC content than introns, which increases DNA bendability^13–16^. In addition, Py tracts (mostly made of Ts) upstream of exons may form a nucleosome energetic barrier^13–16^. Nucleosomes influence splicing by slowing down RNAPII in the vicinity of exons and by modulating the local recruitment of splicing regulators^10,11^. Indeed, depending on their specific chemical modifications (e.g., methylation), histone tails can interact directly or indirectly with splicing factors^17^. Therefore, exon recognition during the splicing process depends on a complex interplay between signals at the DNA level (e.g., nucleosome positioning) and signals at the RNA level (e.g., splicing factor binding sites).

Genes are not randomly organized across a genome, and nucleotide composition bias over genomic regions of varying lengths plays an important role in genome organization at multiple genomic scales. For example, isochores are large genomic regions (≥30 Kbps) with a uniform GC content that differs from adjacent region^18–20^. Isochores can be classified into five families, ranging from less than 37% of GC content to more than 53%^18–20^. GC-rich isochores have a higher density of genes than AT-rich isochores, and genes in GC-rich isochores contain smaller introns than genes in AT-rich isochores^21–23^. It has been proposed that splicing of short introns in a GC-rich context may occur through the intron definition model, while the splicing of large introns in an AT-rich context may occur through the exon definition model^10,24^. Collectively, these observations support a model in which the gene architecture (e.g., size of introns) and gene nucleotide composition bias (e.g., GC or AT content) influence local processes at the exon level, such as nucleosome positioning and intron removal. As exon recognition also depends on the binding to the pre-mRNAs of splicing factors that interact with compositionally-biased sequences, one interesting possibility is that the nature of these splicing factors depends at least in part on the gene nucleotide composition bias. In this setting, we have recently reported that exons regulated by different splicing factors have different nucleotide composition bias^25^.

Here, we have investigated the relationship between the splicing process, gene nucleotide composition bias and chromatin organization at both the local and global levels. We initially identified sets of exons activated by different splicing factors and then demonstrated that analysing the nucleotide composition bias provided a better understanding of the interplay between chromatin organization and splicing-related features, which collectively affect exon recognition. We propose that nucleotide composition bias not only contributes to the 1D and 3D genome organization, but has also local consequences at the exon level during the splicing process.

## Results

### Splicing factor–dependent GC-rich and AT-rich exons

Publicly available RNA-seq datasets generated after knocking down or over-expressing individual splicing factors across different cell lines were analysed (Supplementary Table 1). Using our recently published FARLINE pipeline, which allows the inclusion rate of exons from RNA-seq datasets to be quantified^26^, we defined the sets of exons whose inclusion is activated by each of the 33 splicing factors that were analysed (Supplementary Table 2). We focused on splicing factor–activated exons to uncover the splicing-related features characterizing exons whose recognition depends on at least one splicing factor. We identified 10,707 exons that were activated by at least one splicing factor from the 93,680 exons whose inclusion rate was quantified by FARLINE across all datasets (see Methods).

As expected, splicing factor–activated exons had weaker 3’- and/or 5’-splicing site (ss) scores as compared to the median score of human exons (Supplementary Fig. 1). We computed the nucleotide composition of each splicing factor–activated exon (Supplementary Fig. 2). Note that we systematically refer to both thymine and uracil as “T”, to simplify our goal of analysing sequence-dependent features at both the DNA and RNA levels. In addition, values obtained from splicing factor activated exons were normalized by the median values measured for human coding exons used as a set of control exons, in order to represent results in a consistent way. Sets of exons activated by different splicing factors had a different proportion of GCs as compared to the median values of control exons (Fig. 1a, Supplementary Fig. 3a). Interestingly, the GC content of splicing factor–activated exons positively correlated with the GC content of their flanking introns (Fig. 1b). Accordingly, splicing factor–activated GC- and AT-rich exons were flanked by GC- and AT-rich intronic sequences, respectively (Supplementary Fig. 3b, c). This result is in agreement with previous observations^27^.

**Fig. 1.**
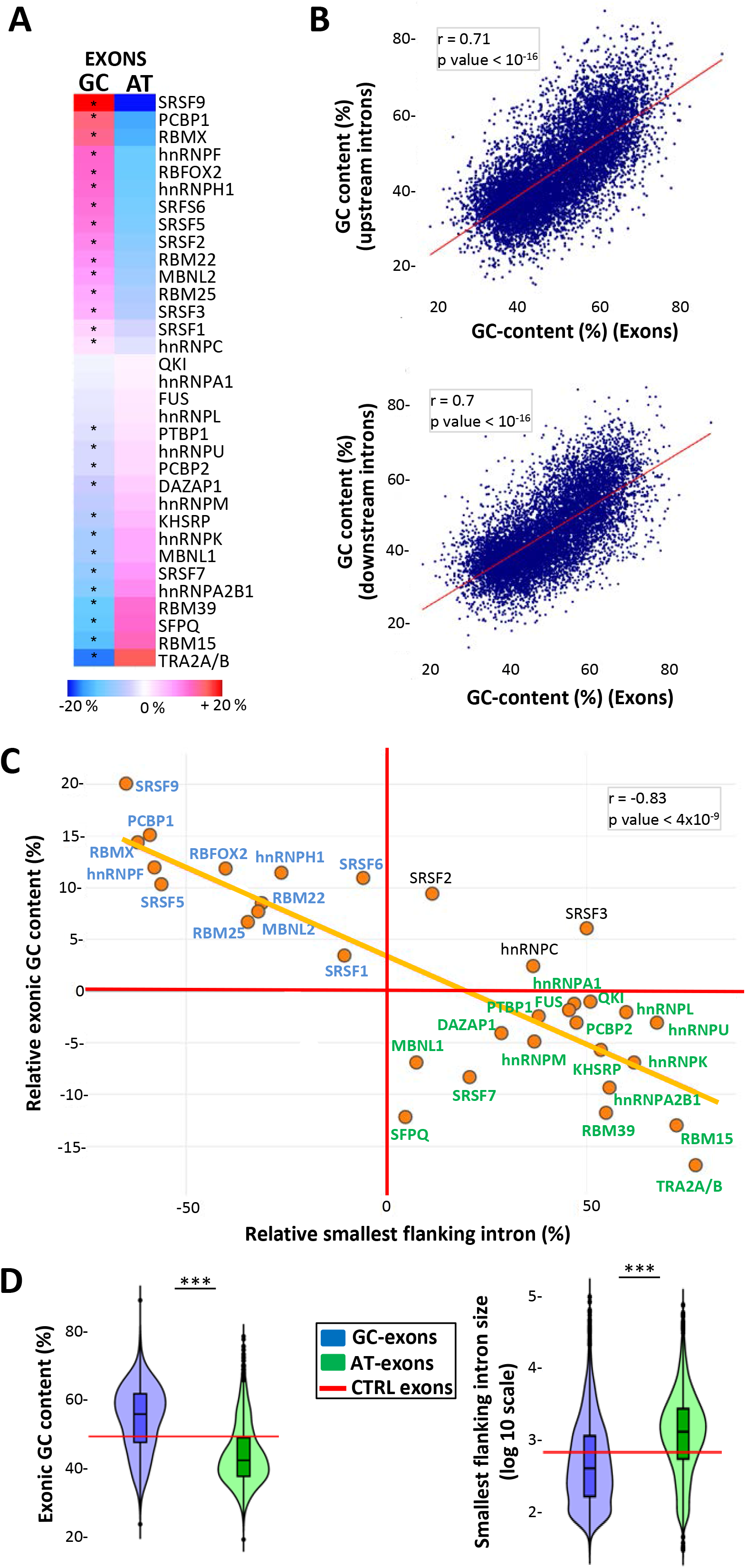
**a** Heatmaps representing the relative median frequency of GC and AT nucleotides in sets of splicing factor–activated exons, as compared to the median values computed from control exons. (*) Student’s test FDR < 0.05. **b** Correlation between the GC content of splicing factor–activated exons and the GC content of their upstream (upper panel) or downstream (lower panel) intron; r = Pearson correlation coefficient. **c** The x-axis represents the relative median size of the smallest intron flanking splicing factor–activated exons, as compared to the median size of human introns. The y-axis represents the relative median GC content of splicing factor–activated exons, as compared to the median GC-frequency of control exons; r = Pearson correlation coefficient. **d** Violin plots representing the GC content (%) of GC-exons and AT-exons (left panel), and the logarithmic nucleotide size of the smallest intron flanking GC-exons and AT-exons (right panel). The red lines indicate the median values computed for control exons. (***) correspond to Wilcoxon’s test *P* < 10^−16^ when comparing GC-exons to AT-exons.

Size analysis of introns flanking splicing factor–activated exons revealed that different sets of splicing factor activated exons were flanked by introns that were either smaller or larger than the median size of human introns (Supplementary Fig. 4a, b). Interestingly, high GC-content was associated with small intron size (Supplementary Fig. 4c), as previously reported^23^. Based on these observations, we defined two groups of exons. The GC-exon group depends on splicing factors activating GC-rich exons that are flanked by small introns (Fig. 1c, in blue), while the AT-exon group depends on splicing factors activating AT-rich exons that are flanked by large introns (Fig. 1c, in green). We excluded for further analyses, exons regulated by SRSF2, SRSF3 or hnRNPC, as these splicing factors regulate GC-rich exons flanked by relatively large introns, as well as exons belonging to both groups (see Materials and Methods). We next analysed different splicing-related features by comparing 3,182 GC-exons to 4,045 AT-exons, representing two populations of exons that: i) differ in terms of both GC-content and flanking intron size, and ii) are activated by distinct splicing factors (Fig. 1c, d, Supplementary Table 2).

### Nucleotide composition bias and splicing-related features

We found that exons and their flanking intronic sequences had similar nucleotide composition biases when considering both whole intronic sequences (Fig. 1b) or intronic sequences located just upstream or downstream exons (Fig. 2a-c). For example, 25 or 100 nucleotide-long intronic sequences that flank GC-exons had a higher frequency of G and/or C nucleotides as compared to intronic sequences flanking AT-exons (Fig. 2a–c). A higher GC-content was associated with a lower minimum free energy measured in a 50 nucleotide-long window centred at the 5’ ss, when comparing GC-exons to both control exons and AT-exons (Fig. 2d, left panel). This suggests a higher stability of base pairing between complementary sequences, and that the 5’ ss of GC-exons are more likely to be embedded in stable secondary structures than AT-exons. A similar feature was observed at the 3’ ss of GC-exons when compared to AT-exons (Fig. 2d, right panel).

**Fig. 2.**
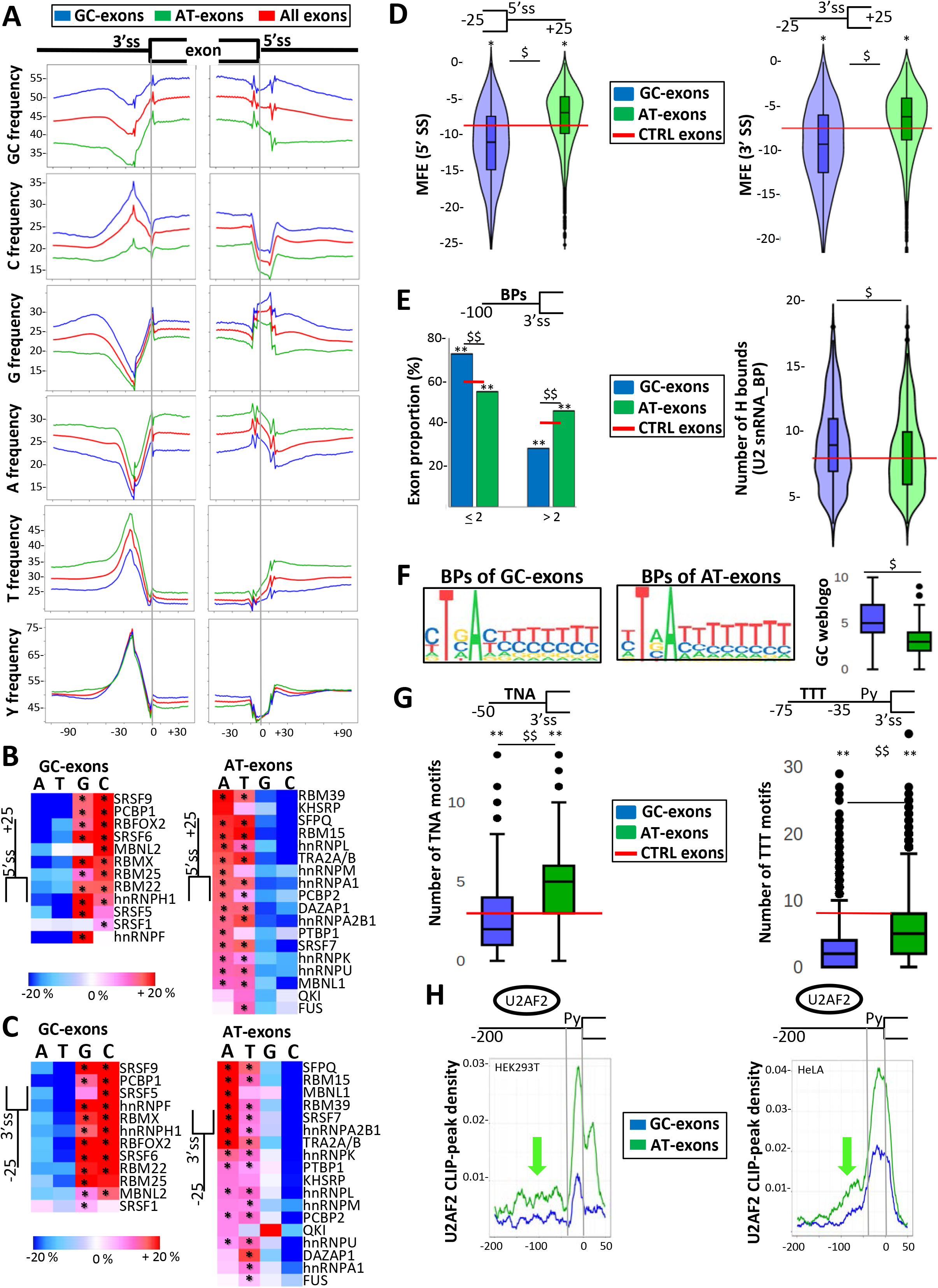
**a** Nucleotide frequency (%) maps in different sets of exons and their flanking intronic sequences. **b** Heatmap representing the average frequency (%, as compared to control exons) of A, T, G or C nucleotides in a window of 25 nucleotides downstream of GC-exons (left panel) or AT-exons (right panel). (*) Wald’s test FDR < 0.05. **c** Heatmap representing the average frequency (%, as compared to control exons) of A, T, G or C nucleotides in a window of 25 nucleotides upstream of GC-exons (left panel) or AT-exons (right panel). (*) Wald’s test FDR < 0.05. **d** Minimum free energy (MFE) at the 5’ ss (left panel) and the 3’ ss (right panel) of GC-exons or AT-exons. MFE was computed using 25 nucleotides within exons and 25 nucleotides within introns. The red lines indicate the median values calculated for control exons. ($) and (*) correspond to Tukey’s test FDR < 10^−16^ when comparing GC-exons to AT-exons, or when comparing GC-exons or AT-exons to control exons, respectively. **e** Proportion (%) of GC-exons or AT-exons with at least two or more predicted BPs in a window of 100 nucleotides in their upstream intron (left panel). Number of hydrogen bonds measured between the U2 snRNA and the BP sequence found in the 25 nucleotides upstream of GC-exons and AT-exons (right panel). The red lines indicate the median values calculated for control exons. (**) and ($$) correspond to Chi^2^ test *P* < 10^−13^ when comparing GC-exons to AT-exons, or when comparing GC-exons or AT-exons to control exons, respectively. ($)Tukey’s test *P* < 0.02 when comparing GC-exons to AT-exons. **f** Weblogos generated using sequences flanking the BPs with the best score in a 25 nucleotide-long window upstream of GC-exons or AT-exons and the boxplot resuming their GC content. ($)Tukey’s test FDR < 10^−16^. **g** Boxplot representing the number of TNA sequences within the last 50 nucleotides of the upstream introns of GC-exons and AT-exons (left panel). Boxplot representing the number of T-rich low complexity sequences in a window between positions −35 and −75 upstream the 3’ss of GC-exons and AT-exons (right panel). The red lines indicate the median values calculated for control exons. ($$) and (**) correspond to Tukey’s FDR < 10^−16^ when comparing GC-exons to AT-exons, and when comparing GC-exons or AT-exons to control exons, respectively. **h** Density of reads obtained from publicly available U2AF2-CLIP datasets generated from HEK293T (left panel) or HeLa (right panel) cells and mapped upstream of GC-exons and AT-exons. The green arrows indicate reads that mapped upstream of the Py tract.

GC-exons were impoverished in Ts but enriched in Cs just upstream of their 3’ ss as compared to control exons (Fig. 2a, c). Therefore, the Py tract of GC-exons was similar to that of control exons (Fig. 2a). On the other hand, AT-exons had a higher frequency of As upstream of their 3’ ss as compared to GC-exons or control exons (Fig. 2a, c). We tested whether the higher frequency of As upstream of AT-exons was associated with a larger number of potential BP sites, which often contain As^28,29^. Indeed, a higher proportion of AT-exons had more than two predicted BPs in their upstream intronic sequence, as compared to GC-exons or control exons (Fig. 2e, left panel). Further, predicted BPs upstream of GC-exons were embedded in sequences that contained a slightly higher proportion of Cs as compared to AT-exons (Fig. 2f). The interaction between the BP and U2 snRNA was reported to be more stable when the BP is embedded in GC-rich sequences^28–30^. Accordingly, the number of hydrogen bonds between BP sites and U2 snRNA was higher for GC-exons than for AT-exons (Fig. 2e, right panel).

As there was a higher frequency of As and Ts upstream of AT-exons as compared to control exons (Fig. 2a, c), we investigated whether this may interfere with the number of potential binding motifs for SF1 (which binds to UNA motifs) and U2AF2 (which binds to U-rich motifs)^1^. As shown in Fig. 2g (left panel), AT-exons contained a larger number of TNA motifs upstream of their 3’ ss as compared to GC-exons. In addition, AT-exons contained a larger number of low-complexity sequences made of three Ts within a four-nucleotide window upstream of the Py tract as compared to GC-exons (Fig. 2g, right panel). Supporting the biological relevance of this observation, the analysis of U2AF2 CLIP-seq datasets revealed a higher U2AF2-related signal upstream of AT-exons, as compared to GC-exons (Fig. 2h), which extended upstream of the Py tract of AT-exons (green arrows), following the pattern of T frequency (Fig. 2a).

### Nucleotide composition bias and dependency for specific spliceosome components

To investigate the interplay between nucleotide composition bias and the dependency of exons on specific spliceosome-associated factors, we analysed publicly available RNA-seq datasets generated after knocking down a variety of spliceosome-associated factors (Supplementary Tables 1 and 2). Exons that were skipped after depletion of SNRPC or SNRNP70 (two components of the U1 snRNP) were in a more GC-rich environment as compared to control exons (Fig. 3a). Likewise, exons that were skipped upon the depletion of the DDX5 and DDX17 RNA helicases (DDX5/17), which enhance exon inclusion by favouring U1 snRNP binding to highly structured 5’ ss^31^, were in a GC-rich environment (Fig. 3a). In addition, the 5’ ss of exons dependent on SNRPC, SNRNP70 or DDX5/17 were predicted to be embedded in stable secondary structures as compared to control exons (Fig. 3b).

**Fig. 3.**
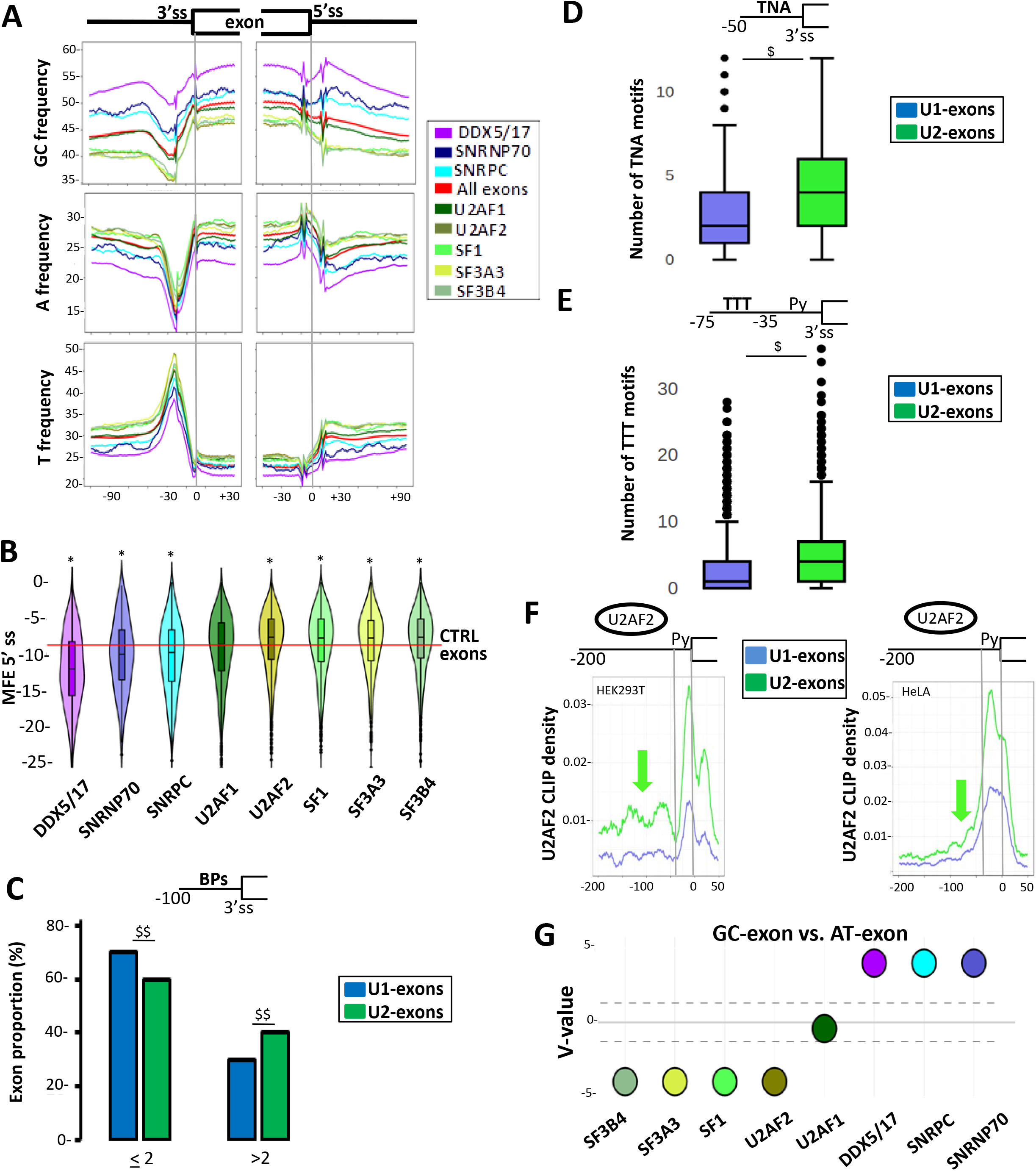
**a** Nucleotide frequency (%) maps in different sets of exons and their flanking intronic sequences. **b** Minimum free energy (MFE) at the 5’ ss of sets of exons activated by different spliceosome-associated factors. MFEs were computed using 25 nucleotides within the exons and 25 nucleotides within the intron. The red line indicates the median values calculated for control exons. (*) Student’s test FDR < 0.02 when comparing a set of exons activated by a spliceosome-associated factor to control (CTRL) exons. **c** Proportion (%) of exons activated by U1 snRNP–associated factors (U1-exons) or by U2 snRNP-associated factors (U2-exons) with two or more predicted BPs in a window corresponding to the last 100 nucleotides in their upstream intron. ($$) corresponds to Chi^2^ test *P* < 10^−16^ when comparing U1-exons to U2-exons. **d** Boxplot of the number of TNA sequences in the last 50 nucleotides upstream introns of U1-exons and U2-exons. ($) corresponds to Wald’s test p-value < 10^−16^ when comparing U1-exons to U2-exons. **e** Boxplot of the number of T-rich low complexity sequences in a window between positions −35 and – 75 upstream the 3’ ss of U1-exons and U2-exons. ($) correspond to Wald’s test p-value < 10^−16^ when comparing U1-exons to U2-exons. **f** Density of reads obtained from publicly available U2AF2-CLIP datasets generated from HEK293T (left panel) or HeLa (right panel) cells and mapped upstream of U1-exons or U2-exons. The green arrows indicate reads mapping upstream of the Py tract. **g** The V-value is a representation of a *P* value calculated by comparing the proportion of GC-exons and AT-exons activated by individual spliceosome-associated factors (see Materials and Methods). A v-value above the dotted line that corresponds to log10 (0.05) is statically significant.

Exons skipped after depletion of SF1, U2AF2, SF3A3 or SF3B4 (but not U2AF1) that recognize splicing signals at 3’-ends of introns were in an AT-rich environment as compared to control exons (Fig. 3a). In addition, a larger proportion of U2-exons—that is, those activated by the U2 snRNP associated factors including SF1, U2AF2, SF3A3 or SF3B4—contained more than two predicted BPs in their upstream intron as compared to U1-exons (e.g., those activated by SNRPC, SNRNP70 and/or DDX5/17) (Fig. 3c). In addition, U2-exons contained a larger number of SF1- and U2AF2-binding sites in their upstream intron as compared to U1-exons (Fig. 3d, e). In agreement with the T-frequency pattern (Fig. 3a), a broader U2AF2-derived signal was observed when comparing U2-exons to U1-exons (Fig. 3f).

To summarize, GC-exons were predicted to have stable secondary structures at their 5’ ss (Fig. 2a, d), and the nucleotide composition bias and splicing-related features of U1-exons were similar to those of GC-exons (Fig. 3a, b). Additionally, the increased frequency of As and Ts upstream of AT-exons (Fig. 2a) was associated with increased numbers of potential decoy signals, and the nucleotide composition bias and splicing-related features of U2-exons were similar to those of AT-exons (Fig. 3a, c–f). We therefore hypothesized that GC-exons were more sensitive to U1 snRNP-than to U2 snRNP-associated factors, in contrast to AT-exons. Accordingly, GC-exons were more likely to be affected by SNRPC, SNRNP70 or DDX5/17 depletion than AT-exons, which were more likely to be affected by SF1, U2AF2, SF3A3 or SF3B4 depletion (Fig. 3g).

### Nucleotide composition bias, gene features and chromatin organization

As exons and their flanking intronic sequences have similar nucleotide composition biases (see Fig. 1b, Fig. 2a, c), we investigated whether the observed local nucleotide composition bias could be extended to the gene level. Indeed, there was a positive correlation between the GC content of exons and the GC content of their hosting gene (Fig. 4a), and GC-exons and AT-exons belong to GC- and AT-rich genes, respectively, as compared to all human genes (Fig. 4b, left panel). In addition, a negative correlation between the GC content of genes and the size of their introns was observed (Supplementary Fig. 4c), as previously reported^23^. Accordingly, GC-exons that are flanked by small introns (Fig. 1) belong to genes containing small introns (Fig. 4b, middle panel). Meanwhile, AT-exons that are flanked by large introns (Fig. 1) belong to large genes containing large introns, as compared to all human genes (Fig. 4b, middle and right panels).

**Fig. 4.**
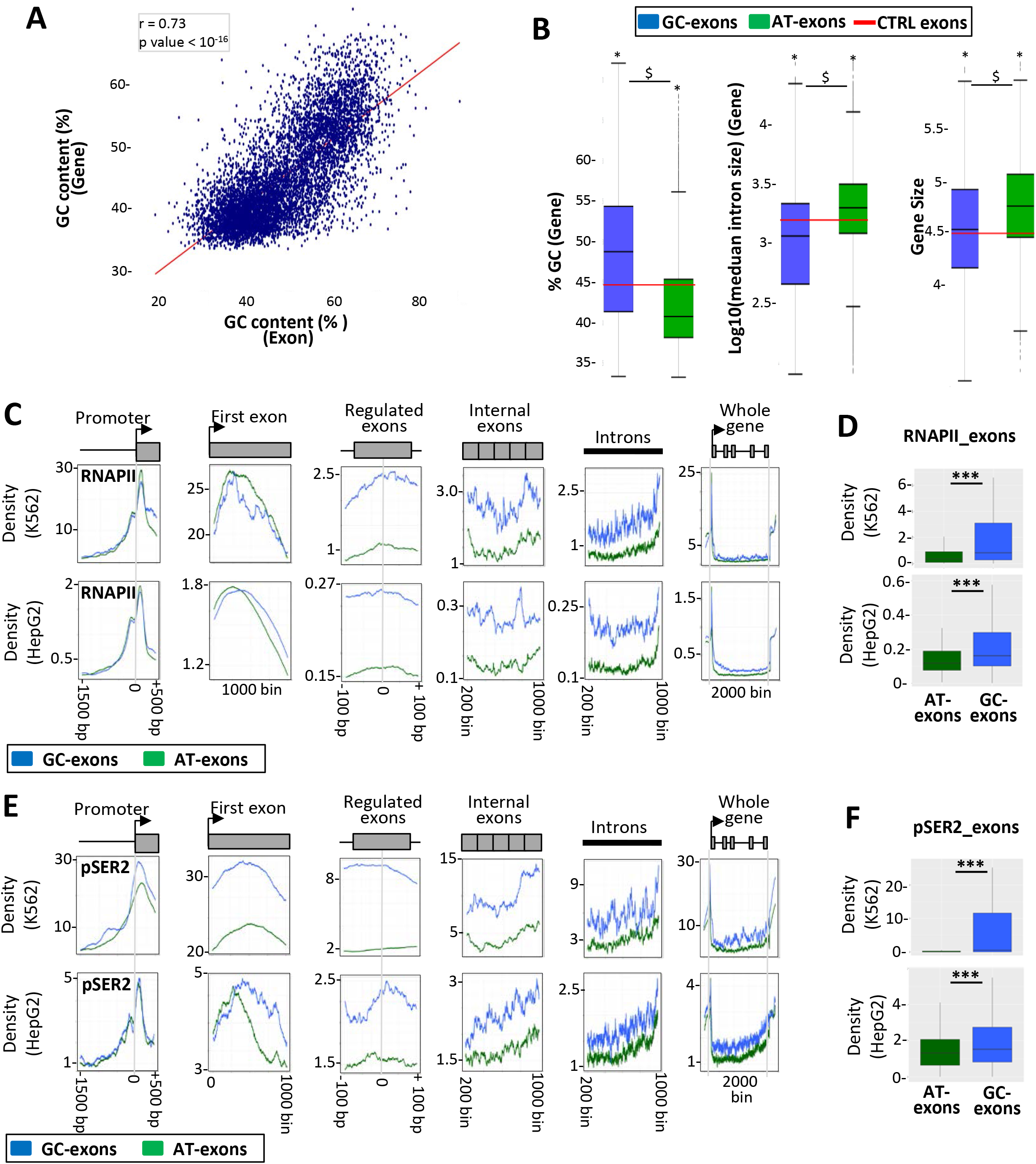
**a** Correlation between the GC content of splicing factor–activated exons and the GC content of their hosting genes; r = Pearson correlation coefficient. **b** Box plots representing the GC content (%, left panel), the intron size (middle panel) and the size (right panel) of genes hosting GC-exons or AT-exons. The red lines indicate the median values calculated for control exons. ($) and (*) correspond to Student’s test (left panel and right panel) and wilcoxon’s test (middle panel) p-value < 2 × 10^−4^ when comparing genes hosting GC-exons and AT-exons, and when comparing genes hosting GC-exons or AT-exons to control genes, respectively. **c** Density of reads obtained after immunoprecipitation of RNAPII in K562 and HepG2 cell lines and then mapped to different parts of genes with GC-exons or AT-exons. **d** Box plots of the mean coverage by RNAPII of GC-exons and AT-exons. (***) Wilcoxon’s test *P* < 10^−6^. e Density of reads obtained after immunoprecipitation of RNAPII phosphorylated at serine 2 (RNAPII-ser2) in K562 and HepG2 cell lines and then mapped to different parts of genes with GC-exons or AT-exons. **f** Box plots of the mean coverage of phosphorylated RNAPII at GC-exons or AT-exons. (***) Wilcoxon’s test *P* < 10^−6^.

In this setting, it has been reported that GC-rich genes are more expressed as compared to AT-rich genes^23,32–34^. Remarkably, for genes comprising either AT-exons or GC-exons, the RNAPII density was similar at promoters and the first exon but was higher at exons and introns of genes hosting GC-exons than genes hosting AT-exons (Fig. 4c, d, Supplementary Fig. 5a). A similar result was observed for the pattern of RNAPII phosphorylated at Ser2 (Fig. 4e, f), suggesting that the RNAPII content on genes hosting GC-exons was likely to be productive.

In addition to be associated with gene expression level, the GC content is associated with nucleosome positioning (see Introduction). The analysis of MNase-seq and ChIP-Seq against H3 datasets across different cell lines revealed a higher nucleosome-density signal on GC-exons than AT-exons (Fig. 5a, Supplementary Fig. 5b), in agreement with a previous report showing that exons embedded in a GC-rich environment have a higher nucleosome density^10,24^. While similar nucleosome-density signals were observed on the first exon of genes hosting either GC-exons or AT-exons, higher signals were observed across all internal exons of genes hosting GC-exons when compared to genes hosting AT-exons (Fig. 5b, c, Supplementary Fig. 5c). In addition, a stronger signal was observed across introns of genes hosting GC-exons compared to genes hosting AT-exons (Fig. 5b, c), in particular across introns flanking splicing factor-activated GC-exons (Fig. 5a). This could be due to the higher frequency of GCs in these introns and the lower frequency of Ts at their 3’-ends when compared to introns flanking AT-exons (Fig. 2). In this setting, there was a marked nucleosome-free region both upstream and downstream of AT-exons when compared to GC-exons (Fig. 5a, green arrows).

**Fig. 5.**
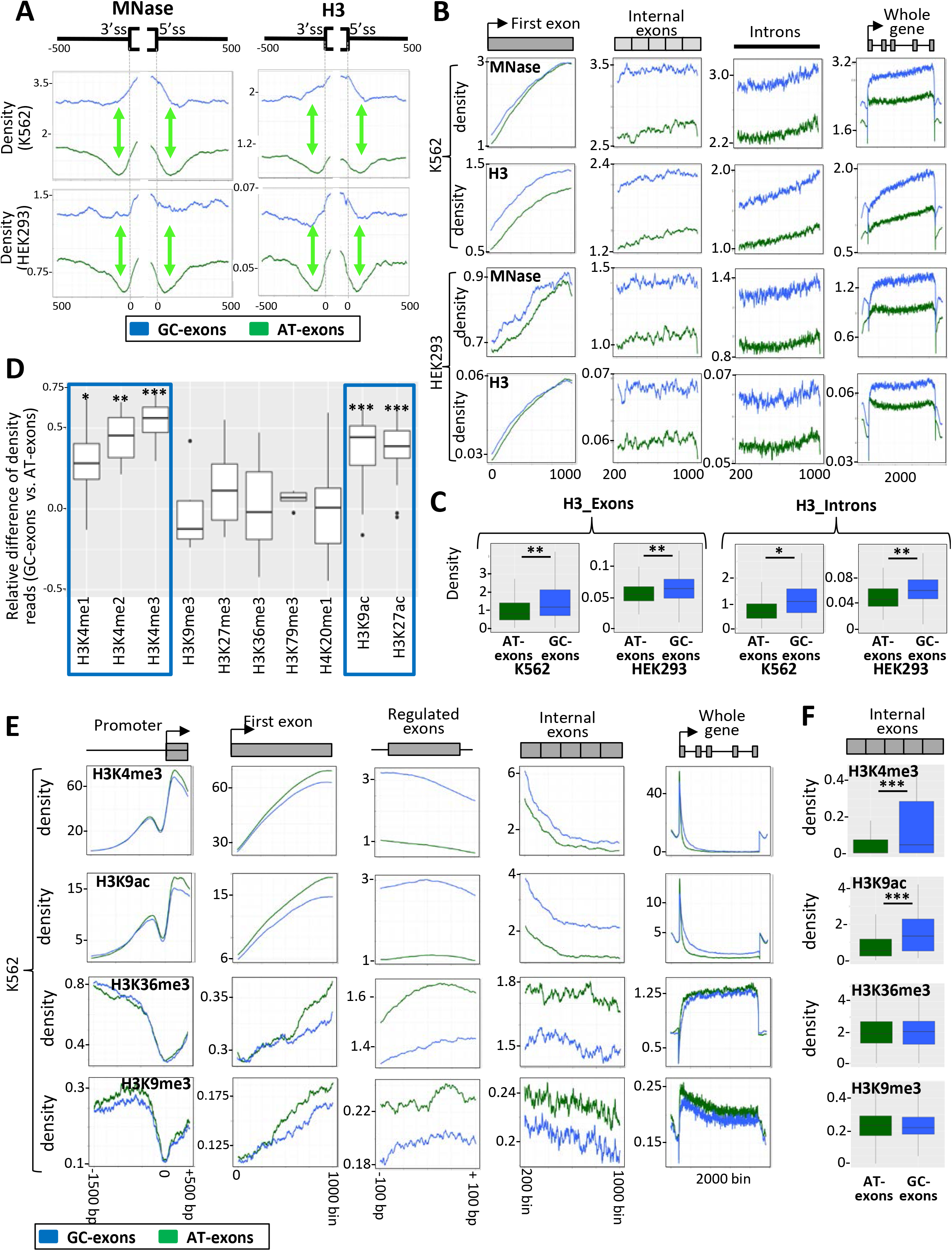
**a** Density of reads obtained after DNA treatment with MNase (left panels), or after immunoprecipitation of the H3 histone (right panels), in K562 and HEK293 cell lines and then mapped to GC-exons or AT-exons and their flanking introns. **b** Density of reads obtained after DNA treatment with MNase, or after immunoprecipitation of the histone H3, in K562 and HEK293 cell lines and then mapped to different parts of genes with GC-exons or AT-exons. **c** Box plots of the mean coverage of reads obtained after immunoprecipitation of the histone H3 from K562 or HEK293 cell lines and then mapped to exons and introns of genes hosting GC-exons or AT-exons. (*) Wilcoxon’s test *P* < 0.05; (**) Wilcoxon’s test *P* <0.001. **d** Box plots representing the relative difference of density reads obtained after DNA immunoprecipitation using antibodies against different histone modifications (as indicated) and then mapped to GC-exons or AT-exons. Each box plot represents the values obtained from several publicly available datasets. (*) Wilcoxon’s test *P* < 0.005; (**) Wilcoxon’s test *P* < 0.001; (***) Wilcoxon’s test *P* < 0.0001. **e** Density of reads obtained from the K562 cell line after immunoprecipitation of DNA using antibodies against different histone modifications (as indicated) and then mapped to different parts of genes with GC-exons or AT-exons. **f** Box plots of the mean coverage of reads obtained after DNA immunoprecipitation using antibodies against H3K4me3, H3K9ac, H3K36me3 or H3K9me3, and then mapped to exons of genes with GC-exons or AT-exons. (***) Wilcoxon’s test *P* < 10^−6^.

The pattern of nucleosomes on GC-exons and AT-exons prompted us to analyze histone tail modifications that play a role in splicing regulation (see Introduction). We analyzed publicly available ChIP-seq datasets generated across different cell lines (Supplementary Table 1). As shown in Fig. 5d, a higher density signal corresponding to H3K4me1, H3K4me2, H3K4me3, H3K9ac, and H3K27ac was detected on GC-exons when compared to AT-exons. No significant differences were observed for H3K9me3, H3K27me3, H3K36me3, H3K79me3, and H4K20me1 (Fig. 5d). The pattern of histone modifications did not seem to be specific to splicing factor-regulated exons. While there was a similar signal density of histone marks on the promoter and first exon of genes hosting either AT-exons or GC-exons, the H3K4me3 and H3K9ac density signals were higher across all the exons of the GC-exons hosting genes (Fig. 5e, f, Supplementary Fig. 5d). Therefore, GC-exons and AT-exons are hosted by genes that have different nucleotide composition bias (i.e., GC content) and architectures (i.e., intron size) and that are embedded in different chromatin environments.

### GC- and AT-exons are hosted by genes belonging to different isochores and chromatin domains

Since GC-exons and AT-exons are hosted by GC- and AT-rich genes, respectively (Fig. 4a, b), and since genes belong to genomic regions (or isochores) having homogenous GC content (see Introduction), we investigated whether there was a correlation between the GC content of exons and the GC content of the isochore they belong to. As shown in Fig. 6a, there was a positive correlation between the GC content of exons and the GC content of their hosting isochores. Accordingly, a larger proportion of GC-exons (>60%) than AT-exons (<25%) belongs to GC-rich isochores (>46% of GC; Fig. 6b). Furthermore, GC-exons and AT-exons cluster in different isochores. Indeed, some isochores contain a larger number of GC-exons than AT-exons, while other isochores contain a larger number of AT-exons (Fig. 6c). This result was confirmed using different annotations of isochores computed with different programs (Supplementary Fig. 6a, b).

**Fig. 6.**
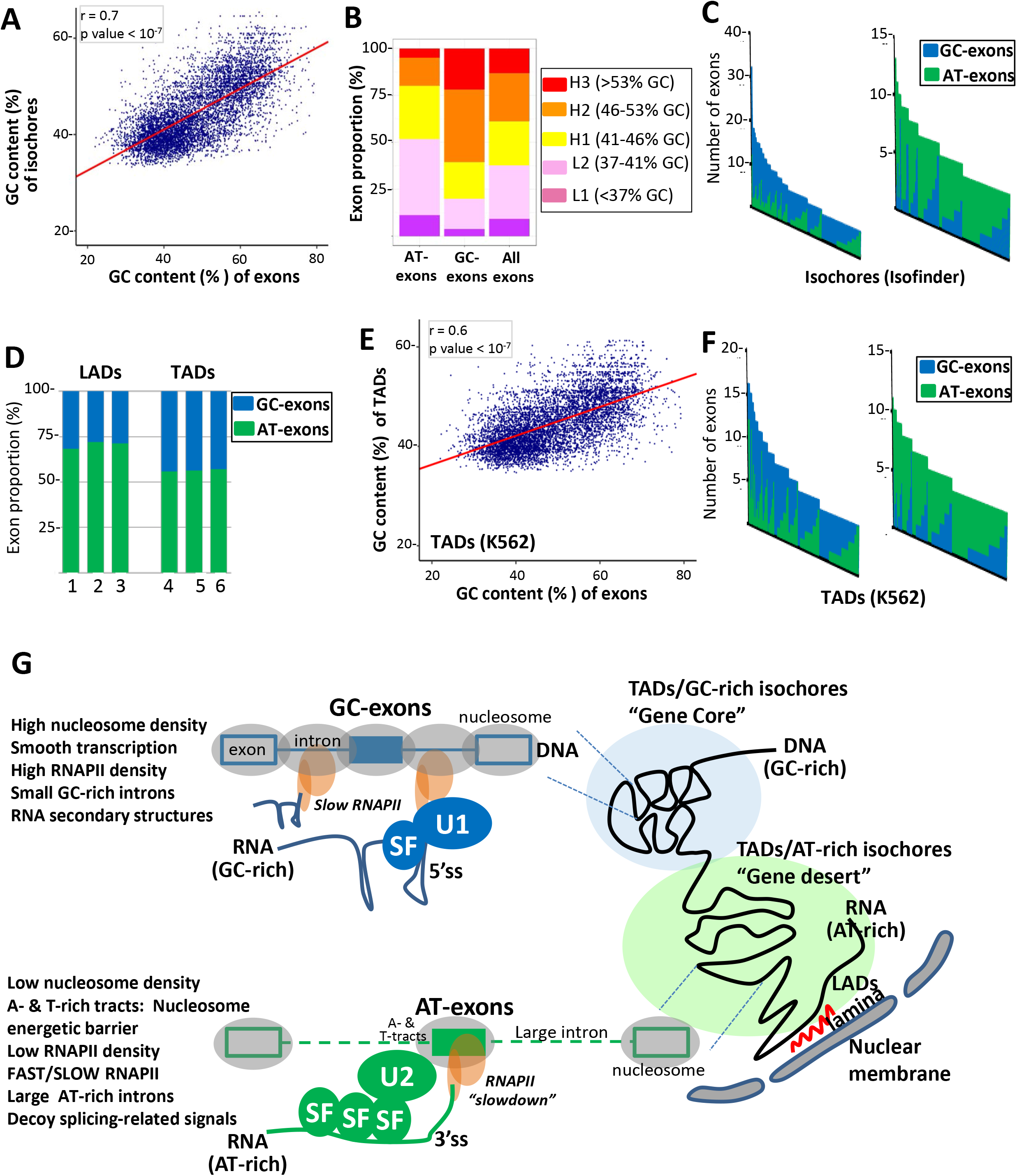
**a** Correlation between the GC content of GC-exons and AT-exons, and the GC content of their hosting isochores defined by ISOFINDER. **b** Proportion of AT-exons, GC-exons and control exons distributed across different isochore families. **c** Number of AT-exons and GC-exons present in individual isochores. Only isochores containing at least five GC-exons or five AT-exons are represented. The left and right panels represent isochores containing preferentially GC-exons or AT-exons, respectively. **d** Proportion of AT-exons and GC-exons in LADs annotated from three different datasets (1, fibroblasts; 2, resting Jurkat cells; 3, activated Jurkat cells) and in TADs annotated from three different cell lines (4, K562; 5, IMR90; 6, MCF7). **e** Correlation between the GC content of GC-exons and AT-exons, and the GC content of their hosting TADs, defined in the K562 cell line. **f** Number of AT-exons and GC-exons present in individual TADs annotated from the K562 cell line. Only TADs containing at least five GC-exons or five AT-exons are represented. The left and right panels represent TADs containing preferentially GC-exons or AT-exons, respectively. **g** GC-rich isochores and TADs contain a large number of genes (“gene core”) that are GC-rich and contain small introns. In contrast, AT-rich isochores, TADs and LADs contain a small number of genes (“gene desert”) that are AT-rich and contain large introns. The regional nucleotide composition bias (over dozens of kbps) increases the probability of local nucleotide composition bias (e.g., at the gene and exon levels). Local nucleotide composition bias influences local chromatin organization at the DNA level (e.g., nucleosome density and positioning) as well as the splicing process at the RNA level. The high density of nucleosomes and GC nucleotides (upper panel) could generate a “smooth” transcription across small genes, favouring synchronization between transcription and splicing. The high density of GC nucleotides increases the probability of secondary structures at the 5’ ss, with consequences on splicing recognition during the splicing process. This constraint could be alleviated by splicing factor (SF; in blue) binding to GC-rich sequences, which enhances U1 snRNP recruitment. The high density of AT nucleotide (lower panel) could favour a sharp difference between exon and intron in terms of nucleotide composition bias, which would favour nucleosome positioning on exons. A- or T-rich sequences located upstream of AT-exons, as well as the present of exonic nucleosomes, could locally (at the exon level) slow down RNAPII, favouring synchronization between transcription and splicing. The high density of AT nucleotides increases the probability of generating decoy signals, such as pseudo BPs or SF1- or U2AF2-binding sites. This constraint could be alleviated by the binding splicing factors (SF, in green) to these decoy signals, thereby enhancing U2 snRNP recruitment.

Some genomic domains, named lamina-associated domains (LADs), are in close proximity to the nuclear envelop^35^ and some DNA domains separated by several dozens of Kbps, named topologically-associated domains (TADs), are in close proximity in the nuclear space^36^. LADs and TADs have been annotated across different cell lines^35,36^. We observed that LADs contain more frequently AT-exons (~70%, Fig. 6d), while TADs contain a similar proportion of GC-exons and AT-exons (~55%). This is in agreement with the fact that LADs correspond to AT-rich regions^35^. Interestingly, we observed a positive correlation between the GC content of exons and the GC content of the TAD they belong to (Fig. 6e). Furthermore, GC-exons and AT-exons cluster in different TADs. Indeed, some TADs contain a larger number of GC-than AT-exons, while other TADs contain a larger number of AT-exons (Fig. 6f). This result was confirmed using different annotations of TADs across different cell lines (Supplementary Fig. 6c, d). Collectively, these observations support a model where nucleotide composition bias establishes a link between genomic organization (e.g., isochores or TADs) and the splicing process.

## Discussion

The rules that govern exon recognition during splicing and that explain the dependency of some exons on different classes of splicing factors remain to be clarified. We propose that nucleotide composition bias (e.g., GC content) over large genomic regions that plays an important role in genome organization, leaves a footprint locally at the exon level and induces constraints during the exon recognition process that can be alleviated by local chromatin organization and different classes of RNA binding proteins.

The human genome is divided in isochores corresponding to regions of varying lengths (up to several dozens of Kbps) having a uniform GC-content that differs from adjacent regions^18–20^. GC-rich isochores have a higher density of genes than AT-rich isochores and GC- and gene-rich genomic regions are highly expressed^19–23,32–34^ (Fig. 6g). Increasing evidence indicates that the one-dimensional genome organization, defined by regional nucleotide composition bias, is related to the three-dimensional genome architecture. For example, an overlap between isochores and TADs has been reported^19,37^. Both the relationship between the 1D and 3D genome organization and the higher density of actively transcribed genes in GC-rich regions could be explained by the physicochemical properties of GC and AT nucleotides. For example, the nature of the base stacking interactions between GC nucleotides increases DNA structural polymorphism that in turn increases DNA bendability and flexibility with consequences on DNA foldin^13–15,38–41^. Furthermore, the higher stability of G:C base pairing and frequency of GC-associated polymorphic structures have been proposed to increase the resistance of the DNA polymer to transcription-associated physical constraints. For example, the GC content is associated with the transition from the B-to Z-DNA form, the former “absorbing” the topological and torsional stresses that are generated during transcription^38–42^. In this setting, we observed that GC-exons and their hosting GC-rich genes have a higher RNAPII density than AT-exons and their hosting AT-rich genes (Fig. 4). Based on these observations, transcription and the three-dimensional genome organization may constrain the nucleotide composition of genomic regions, which in turn has “local” consequences on exon recognition during co-transcriptional splicing.

Supporting this possibility, first we observed a positive correlation between the GC content of exons, their flanking introns, and their hosting-genes, -isochores, and -TADs (Fig. 1b, Fig. 4a, Fig. 6a, e), in agreement with the notion that the GC content is uniform and homogenous regardless of the genomic scale^18–20^. Second, differential GC content is associated with specific constraints on the exon recognition process. For example, our analyses and reports from the literature support a model where a high local GC-content favours the formation of RNA secondary structures that can hinder the recognition of the 5’ ss by occluding them, therefore limiting the access of U1 snRNA to the 5’ ss^2–5^. Accordingly, exons sensitive to the depletion of SNRPC and SNRNP70, two components of the U1 snRNP, as well as exons sensitive to the DDX5 and DDX17 helicases that enhance the recognition of structured 5’ ss owing to their RNA helicase activity^31^, are embedded in a GC-rich environment (Fig. 3a, b). In this setting, splicing factors that activate GC-exons (for instance, hnRNPF, hnRNPH, PCBP1, RBFOX2, RBM22, RBM25, RBMX, SRSF1, SRSF5, SRSF6 and SRSF9) bind to G-, C-, or GC-rich motifs^7–9^ (Supplementary Fig. 7). Furthermore, hnRNPF, hnRNPH, RBFOX2, RBM22, RBM25 and several SRSF splicing factors are known to enhance U1 snRNP recruitment^43–48^. Therefore, a high GC-content could increase the probability of generating secondary structure at the 5’ ss, which decreases exon recognition. Simultaneously, high GC-content could increase the recruitment of splicing factor binding to GC-rich motifs, thereby enhancing U1 snRNP recruitment (Fig. 6g). While RNA secondary structures at the 5’ ss negatively impact exon recognition, structures at the 3’ ss favour exon recognition and replace the requirement for U2AF2 in splicing^4,5^. In addition, a high GC-content, as well as G- and C-rich motifs upstream of the BP, enhance U2 snRNA binding and BP recognition^28,29,49^. Accordingly, exons embedded in a GC-rich environment are more sensitive to factors associated with U1 snRNP than those with U2 snRNP (Fig. 3g).

Our results also support a model in which a high content of AT nucleotides in large introns can negatively influence exon recognition. Indeed, high AT-content upstream of exons associated with a larger number of potential BPs (Fig. 2e), in agreement with a previous report^30^. A high AT-content upstream of exons also associated with a larger number of SF1- and U2AF2-binding sites (Fig. 2g, h). In this setting, increasing evidence indicates that binding of spliceosome-associated factors (e.g., SF1 or U2AF2) to pseudo-signals or decoy signals can inhibit splicing by decreasing the efficiency of spliceosome assembly^29,50–55^. These observations suggest that splicing factors that activate AT-exons enhance exon recognition either by enhancing U2 snRNP recruitment or by binding to decoy splicing signals. Accordingly, splicing factors activating AT-exons, including hnRNPA1, hnRNPM, RBM15, RBM39, SFPQ and TRA2 interact with and enhance the recruitment of SF1, U2AF2, U2AF1 and/or U2 snRNP^56–61^. In addition, some of these splicing factors can compete with U2AF2 or SF1 for binding to intronic 3’-end splicing signals, and binding of splicing factors such as hnRNPA1 and PTBP1 to decoy splicing signals has been proposed to “fill up” a surplus of splicing signals and consequently enhancing the recognition of *bona fine* splicing sites^29,50–55^. Therefore, a high AT-content could increase the probability of generating decoy splicing signals at intronic 3’-ends, which would decrease exon recognition. However, a high AT-content could simultaneously increase the probability of recruiting splicing factors at decoy signals, thereby strengthening the recruitment of spliceosome-related components (e.g., SF1 and U2AF2) to *bona fide* splicing signals and ultimately enhancing exon recognition (Fig. 6g).

The synchronization between transcription and splicing plays a major role in the exon recognition process^10,11,62,63^. We propose that the coupling between transcription and splicing operates through different mechanisms depending on the gene nucleotide composition bias, which impacts chromatin organization and RNAPII dynamics. Indeed, at the chromatin level, nucleosomes are better positioned on exons in an AT-rich context than in a GC-rich context, and there is a higher density of nucleosomes in both GC-rich exons and introns (Fig. 5a–c), as already reported^13–16^. This feature could result from the fact that exons embedded in an AT-rich context have a much higher GC content than their flanking intronic sequences, in contrast to exons embedded in a GC-rich context^10,24^ (Fig. 2a). In this setting, GC-rich stretches favour DNA wrapping around nucleosomes, because the stacking interactions between GC nucleotides allow DNA structural polymorphism that in turn increases DNA bendability; in contrast, T- and A-rich stretches form more rigid structures that create nucleosome energetic barriers^13–16^. Therefore, increasing the intronic GC-content increases the probability of nucleosomes sliding from exons to introns, while increasing the density of Ts and As in exonic flanking regions creates nucleosome energetic barriers. Consequently, transcription and splicing synchronization in an AT-rich environment could depend on nucleosomes being well-positioned on exons, as these would locally slow down RNAPII and thereby favour recruitment of splicing-related factors (Fig. 6g), as previously proposed^10^. Of note, components associated with the U2 snRNP interact with chromatin-associated factors^64,65^.

Synchronization between transcription and splicing could rely on other mechanisms when exons are within a GC-rich environment. Indeed, both the higher density of nucleosomes across introns of GC-rich genes (Fig. 5a–c), and the higher stability of G:C basepairing, create constraints that reduce the velocity of RNAPII across both exons and introns. Accordingly, the rate of elongation by RNAPII is negatively correlated with gene GC-content^66^. Therefore, high GC-content may facilitate the synchronization between transcription and splicing by “smoothing” RNAPII dynamics all along GC-rich genes. Of note, extensive interactions between the U1 snRNP and RNAPII-associated complexes have been reported^67^; therefore, a slower speed of RNAPII across GC-rich genes may facilitate U1 snRNP recruitment (Fig. 6g). Further supporting that gene GC-content plays an important role in the interplay between gene expression levels and splicing, intron removal occurs more efficiently in highly-expressed genes^68^, and GC-rich genomic regions associate with nuclear speckles^69,70^.

Altogether, these observations suggest a link between nucleotide composition bias, genome organization and RNA processing (Fig. 6g). We propose a model in which transcription and genome organization constrain the nucleotide composition of DNA over dozens of Kbps. In turn, nucleotide composition bias induces local (at the exon level) constraints on the splicing process by affecting specific splicing-related features. However, constraints induced by nucleotide composition bias can be alleviated by specific mechanisms. For example, although AT-exons are weakened in terms of intron 3’-end definition, this can be alleviated by an interplay between exon-positioned nucleosomes, U2 snRNP-associated factors and splicing factors that bind decoy signals and/or enhance U2 snRNP recruitment. Likewise, while GC-exons are weakened at their 5’ ss because of the formation of RNA secondary structures, this can be alleviated by an interplay between a slow RNAPII, U1 snRNP-associated factors and splicing factors that bind to GC-rich sequences and enhance the recruitment of the U1 snRNP. In this model, splicing factors would enhance the recognition of exons by counteracting splicing-associated constraints resulting from nucleotide composition bias and providing room for regulatory processes such as alternative splicing.

## Materials and Methods

### RNA-seq dataset analyses and establishment of GC- and AT-exon sets

Publicly available RNA-seq datasets generated from different cell lines transfected with siRNAs or shRNAs targeting specific splicing factors, or transfected with splicing factor expression vectors, were recovered from GEO and ENCODE (Supplementary Table 1). RNA-seq datasets were analysed using FARLINE, a computational program dedicated to analyse and quantify alternative splicing variations, as previously reported^26^. This study focused on exons whose inclusion depends on at least one splicing factor. For this, the sets of exons that are activated by each analysed splicing factor in at least one sample were defined (Supplementary Table 2). Exons that are regulated in an opposite way by the same splicing factor in different samples were eliminated. GC-exons were defined as those being activated by splicing factors (i.e., SRSF9, PCBP1, RBMX, hnRNPF, RBFOX2, SRSF5, hnRNPH1, RBM22, RBM25, MBNL2, SRSF6 and SRSF1) that enhance the inclusion of GC-rich exons flanked by small introns. AT-exons were defined as those being activated by splicing factors (i.e., TRA2A/B, RBM15, RBM39, hnRNPA2B1, KHSRP, hnRNPM, SRSF7, SFPQ, MBNL1, DAZAP1, PTBP1, hnRNPL, hnRNPK, FUS, QKI, hnRNPA1, PCBP2 and hnRNPU) that enhance the inclusion of AT-rich exons flanked by large introns. For further analyses, exons found in the two-exon sets (i.e., exons regulated by two splicing factors of different classes) were eliminated (about 25%), leading to one list of 3,182 GC-exons and another of 4,045 AT-exons (Supplementary Table 2). U1-exons were defined as exons activated by the SNRPC, SNRNP70 and/or DDX5/17 factors, and U2-exons were defined as exons activated by the U2AF2, SF3B4, SF1 and/or SF3B4 factors. All genomic annotations were from FasterDB^71^.

### Heatmaps and frequency maps

Heatmaps represent the median value for a given feature of a set of splicing factor activated exons as compared to the median value obtained for control exons. The formula (1) was used to compute the relative value of a feature D in a set of *n* exons *SI* compared to a set of *m* control exons *C.* The control set of exon used corresponds to human exons annotated in FASTERDB.

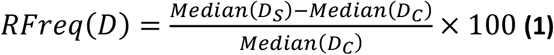

Where, *D_S_* and *D_C_* are the vectors of *D* values for the sets *S* and *C.* Heatmaps of fig 2b and 2c were generated using a *Mean* function in the formula (1). A linear model (with R lm()and summary() functions) was used to compare the GC content of exons activated by each splicing factor to the GC content of the control exons. With this model, a Student’s test was computed between control exons and each set of exons activated by a splicing factor. A generalized linear model for the Poisson distribution (with glm()and summary() functions) was used to compare the proportion of As, Gs, Cs, or Ts, in the 25 first nucleotides upstream exons between exons activated by each splicing factor to control exons. With this model, a Wald’s test was computed between each splicing factors activated set of exons and control exons. The sequences corresponding to the intron-exon junctions (the last 100 nucleotides of the upstream intron and the first 50 nucleotides the exon) were recovered for each exon. The mean frequency of a given nucleotide was computed at each window position using sliding window with a size and a step of 20 and 1 nucleotide, respectively. The same procedure was applied for the sequences corresponding to exon-intron junctions defined as the last 50 nucleotides of the exon and the first 100 nucleotides of the downstream intron.

### Splice site scores, minimum free energy, BP predictions, U2 binding energy and motif count

The splice site scores were calculated for each FasterDB exons with MaxEntScan (http://genes.mit.edu/burgelab/maxent/Xmaxentscan_scoreseq.html). The minimum free energy (MFE) was computed from exon-intron junction sequences (25 nucleotides within the intron and 25 nucleotides within the exon) using RNAFold from the ViennaRNA package (v2.4.1; http://rna.tbi.univie.ac.at/cgi-bin/RNAWebSuite/RNAfold.cgi))’ These sequences were split in two groups: the sequences centered on the 5’ ss and the sequences centered on 3’ ss. The Anscombe transformation was applied on MFE values to obtain Gaussian distributed values. An ANOVA model (with R, aov() function) was built and statistical differences between every couple of group of exons was tested with a Tukey’s test. Differences between MFE of exons activated by each spliceosome associated factor and control exons were tested using a linear model (with R lm()and summary() functions). The number of branch points in a given sequence corresponding to the 100 nucleotides preceding 3’-ss was computed with SVM-BP finder (http://regulatorygenomics.upf.edu/Software/SVM_BP/). Only branch point sites with a svm score > 0 were considered. The U2 snRNA binding energy corresponds to the number of hydrogen bounds between the nucleotides surrounding the branch point of an RNA sequence (without the branchpoint adenine) and the branch point binding sequence of U2 snRNA. The RNAduplex script in the ViennaRNA package (v2.4.1) was used to determine the optimal hybridization structure between the branch point binding sequence of U2 snRNA (GUGUAGUA) and the RNA sequence. The RNA sequence is composed of 5 nucleotides before and 3 after the branch point. Then, the sum of hydrogen bounds forming between the RNA and the U2 sequence were computed. The number of TNA motifs was computed in the last 50 nucleotides of each intron. To test the differences for the three features mentioned above between groups of exons activated by spliceosome-associated factors and the control group of exons, the same procedure as the one applied for fig 2b and 2c was used (see previous section) (fig 3c and 3d). To test the differences between every couple of group of exons, a Tukey’s test was used (with R, glh function (library multcomp)) (fig 2e and 2g, left panel).

T-rich low complexity sequences were computed between the 75^th^ to the 35^th^ nucleotides upstream the 3’ ss, using a sliding window (size 4, step 1 and at least three Ts). Statistical differences were tested by using a linear model for the negative binomial distribution (with glm.nb()function, library MASS). Statistical differences between every couple of group of exons were tested using a Tukey’s test (R, glh function).

### V-value: Exons regulation by U1 or U2 snRNP–associated factors

Difference between the proportion of GC-exons and AT-exons depending on spliceosome-associated factors was tested by a randomization procedure. For each spliceosome-associated factor, 10,000 subsamples of AT-exons (with the size of the GC-exons set) were generated and the proportion of exons activated by the factor for each sample was computed. The empirical p-value *p_emp_* was computed as:

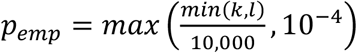

with *k* the number of AT-exon samples with a higher or equal proportion of exons activated by the factor of interest as compared to GC-exons and *l* the number of AT-exons samples with a lower or equal proportion of exons activated by the factor of interest as compared to GC-exons.

The V-value for each spliceosome-associated factor was computing using the formula:

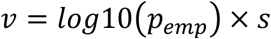

Where *s* = 1 if *k* > *l*; *s* = −1 otherwise.

#### Statistical analysis at gene level

To test whether the GC content of genes hosting AT-exons (AT-genes) or GC-exons (GC-genes) was different, the GC content of the genes according to their size and their group was modeled with an anova model (in R, aov() function). A Tukey’s test on this model was computed to compare between all the possible couples of gene groups (AT-, GC-, and control-genes). To test if the median intron size was different for each couple of AT-, GC- and control groups of genes a Wilcoxon’s test was performed. To test if the gene size of GC-, AT- and control genes was different, we built an anova model (in R, aov() function). Then a Tukey’s test was performed on this model. For those analyzes genes hosting both AT- and GC-exons were not considered.

#### CLIP-seq dataset analyses

Bed files from publicly available CLIP-seq datasets generated using U2AF2 antibodies (GSE83923, GSM2221657; GSE61603 or GSM1509288) were used to generate density maps. The bed files were first sorted and transformed into bedGraph files using the bedtools suite (v2.25.0). The bedGraph files were then converted into bigWig files using bedGraphToBigWig (v4). The 5’ ss and 3’ ss regions (comprising the ss, 200 nucleotides into the intron and 50 nucleotides into the exon) were considered. The proportions of GC- or AT-exons, or exons activated by at least one U1-associated factor (e.g., SNRPC, DDX5/17 and SNRNP70) or at least one U2-associated factor (e.g., U2AF2, SF3B4, SF1 and SF3B4) with CLIP peak signals at each nucleotide position of the 5’ ss and 3’ ss regions, were computed.

#### Analysis of MNase datasets and RNAPII, H3 and histone mark–related ChIP-seq datasets

ChIP-seq or MNase-seq datasets were recovered from Cistrome, ENCODE and GEO databases (Supplementary Table S1). Coverage files (BigWig) were directly downloaded from Cistrome and RNAPII ChIP-seq from ENCODE. Annotations were lifted over from hg38 to hg19 if the coverage file came from Cistrome database. Otherwise, raw data were downloaded for analysis with homemade pipelines. Reads were trimmed and filtered for a minimum length of 25b using Cutadapt 1.16 (options: −m 25), trimmed at their 3’ end for a minimum quality of 20 (−q 20) and then filtered for minimum length of 25b (–m 25). The processed reads were mapped to hg19 with Bowtie2 2.3.3 (options: --very-sensitive --fr −I 100 −X 300 --no-mixed) and filtered for mapping quality over 10 with samtools view 1.6 (options: −b −q 10). For ChIP-seq experiments generated using sonication only, duplicates were removed with homemade tools, which check for coordinates and CIGAR of the Read and the Read 2 adaptor sequence if paired-end sequencing was used. Fragments were reconstituted from the reads, and fragment-coverage files were built using MACS2 2.1.1.20160309 (options: −g hs −B). The metaplots of ChIP/MNase-seq on genes were generated by recovering the fragment coverage (promoter: −1500b/+500 from the TSS; first exon, internal exons and introns: according to the coordinates of the annotation; splicing factor–regulated exons: −100 / +100 from the center, or 500b into the intron and 50b into the exon from the splicing site [MNase and H3]; whole gene: according to the coordinates of the annotations and – 200/+200 from the annotation). In the case of RNAPII coverage, only exons regulated in the corresponding cell line, or annotation from their hosting gene, were considered. For internal exons and introns, the coverages of the annotations from the same gene were concatenated respecting their genomic order. The coverages recovered according to the coordinates of the annotations were the split into 1000 bins. The first 199 bins of “internal exons” or of introns were removed to avoid displaying signals influenced by the promoter. Metaplots were built by computing, at each position or bin, the average coverage across the annotations. Statistics were done by comparing average coverage in the annotations from two groups, which were cell-line specific, with a Wilcoxon’s test. In Fig. 5d, the average of the mean coverage on each regulated exons (−100/+100 from center) was computed. For each ChIP-seq experiment, the ratio of the averages (“GC” – “AT”/max(“GC”, “AT”) was computed and used to build the boxplot in Fig. 5c, and the statistics were computed with a Wilcoxon’s test of the average per experiment of GC-compared to AT-exons.

#### Isochores, LADs, TADs, and CLIP-seq dataset analyses

Chromosome coordinates of isochores, LADs and TADs were recovered from previous publications or GEO (see Supplementary Table 3). The bedtool intersect command was used to determine the isochore, TAD and LAD regions to which each exon belongs. The percentage of GC, and the number of exons (GC- or AT-exons), present in each annotated isochore, LAD or TAD was calculated (Supplementary Table 3).

## Acknowledgments

We gratefully acknowledge the support from the PSMN (Pôle Scientifique de Modélisation Numérique) of the ENS de Lyon for the computing resources. We thank the members of the LBMC Biocomputing Center for their involvement in this project. This work was funded by Fondation ARC (PGA120140200853), INCa (2014-154), ANR (CHROTOPAS), AFM-Téléthon and LNCC. J.B.C was supported by Fondation de France. None of the authors have any competing interests.

