## Supplementary Figures for "Interplay between gene nucleotide composition bias and splicing"

**Supplementary Table S1:** GEO number of publicly available datasets analyzed in this work.

**Supplementary Tables S2:** List of exons activated by each analyzed splicing factors and lists of the GC-exons and AT-exons.

**Supplementary Tables S3:** Annotation of isochores, LADs, and TADs.

**Figure S1**

Violin plots representing the relative 3' ss score (upper panel) and the relative 5' ss score (lower panel) for each set of splicing-factor activated exons, when compared to control exons. CCE=control coding exons; (*) Wilcoxon's test FDR < 0.05.

**Figure S2**

Violin plots representing the relative adenine, cytosine, guanine, and thymine frequencies for each set of splicing-factor activated exons, when compared to control exons. CCE=control coding exons; (*) Student's test FDR < 0.05.

**Figure S3**

**A.** Violin plots representing the relative GC frequency in each set of splicing factor-activated exons, when compared to control exons . CCE=control coding exons; (*) Student's test FDR < 0.05.

**B.** Violin plots representing the relative GC frequencies in introns that are upstream and downstream, respectively, of splicing factor-activated exons. CCE=control coding exons; (*) Wilcoxon's test FDR < 0.05.

**C.** Heatmaps representing the relative frequency of GC and AT nucleotides in the upstream (n-1) and downstream introns of splicing factor-activated exons. (*) Wilcoxon's test, FDR < 0.05.

**Figure S4**

**A.** Violin plots representing the relative upstream intron size (upper panel) and the relative downstream intron size (lower panel) for each set of splicing-factor activated exons, when compared to control exons. CCE=control coding exons

**B** Heatmap representing the median size of the smallest intron flanking splicing factor–activated exons, when compared to the median size of human introns. The sets of splicing factor–activated exons are represented in the same order as in fig 1a.

**C.** Correlation between the relative median size of introns of genes hosting splicing factor-activated exons (compared to all human introns) and the relative gene GC-content (compared to all human genes); r = Pearson correlation coefficient.

**Figure S5**

**A.** Density of reads obtained after immunoprecipitation of RNAPII in HEK293 cell line and mapping to different parts of the genes hosting GC-exons or AT-exons.

**B.** Density of reads obtained after DNA treatment with MNase (left panel) or after immunoprecipitation of the histone H3 (right panel) in Hela cells and mapping to GC-exons or AT-exons and their flanking introns.

**C.** Density of reads obtained after DNA treatment with MNase or after immunoprecipitation of the histone H3 in Hela cells and mapping to different parts of the genes hosting GC-exon or AT-exons.

**D.** Density of reads obtained from the HEK293 and Hela cell lines after immunoprecipitation of DNA using antibodies against H3K4me3 and H3K9ac and mapping different parts of genes hosting GC-exons or AT-exons.

**Supplementary Figure S6**

**A.** Correlation between the GC content of GC-exons and AT-exons and the GC content of their hosting isochores (left panel) defined by *Constantini et al* (Constantini, M., Clay, O., Auletta, F. & Bernardi, G. An isochore map of human chromosomes. 2006; Genome Res 16, 536-541). Proportion of AT-exons, GC-exons, and all human exons distributed across different isochore families defined by *Constantini et al.* (middle panel). Number of AT-exons and GC-exons present in individual isochores defined by *Constantini et al* (right panel). The left and right panels, represent, isochores containing preferentially GC-exons or AT-exons, respectively.

**B.** Same as in Supplementary Fig 6a but using isochores defined by Isosegmenter (https://github.com/bunop/isoSegmenter; Cozzi, P., Milanesi, L. & Bernardi, G. Segmenting the Human Genome into Isochores. 2015; Evolutionary bioinformatics online 11, 253-261).

**C.** Correlation between the GC content of GC-exons and AT-exons and the GC-content of their hosting TADs defined in the IMR90 cell line (left panel). Number of AT- and GC-exons present in individual TADs annotated from the IMR90 cell line (right panel).

**D.** Same as in Supplementary fig 6c but using TADs defined in the MCF7 cell line.

**Supplementary Figure S7**

Splicing factor binding motifs retrieved from different resources. Splicing factors in blue color activate GC-exons, while splicing factors in green color activate AT-exons.

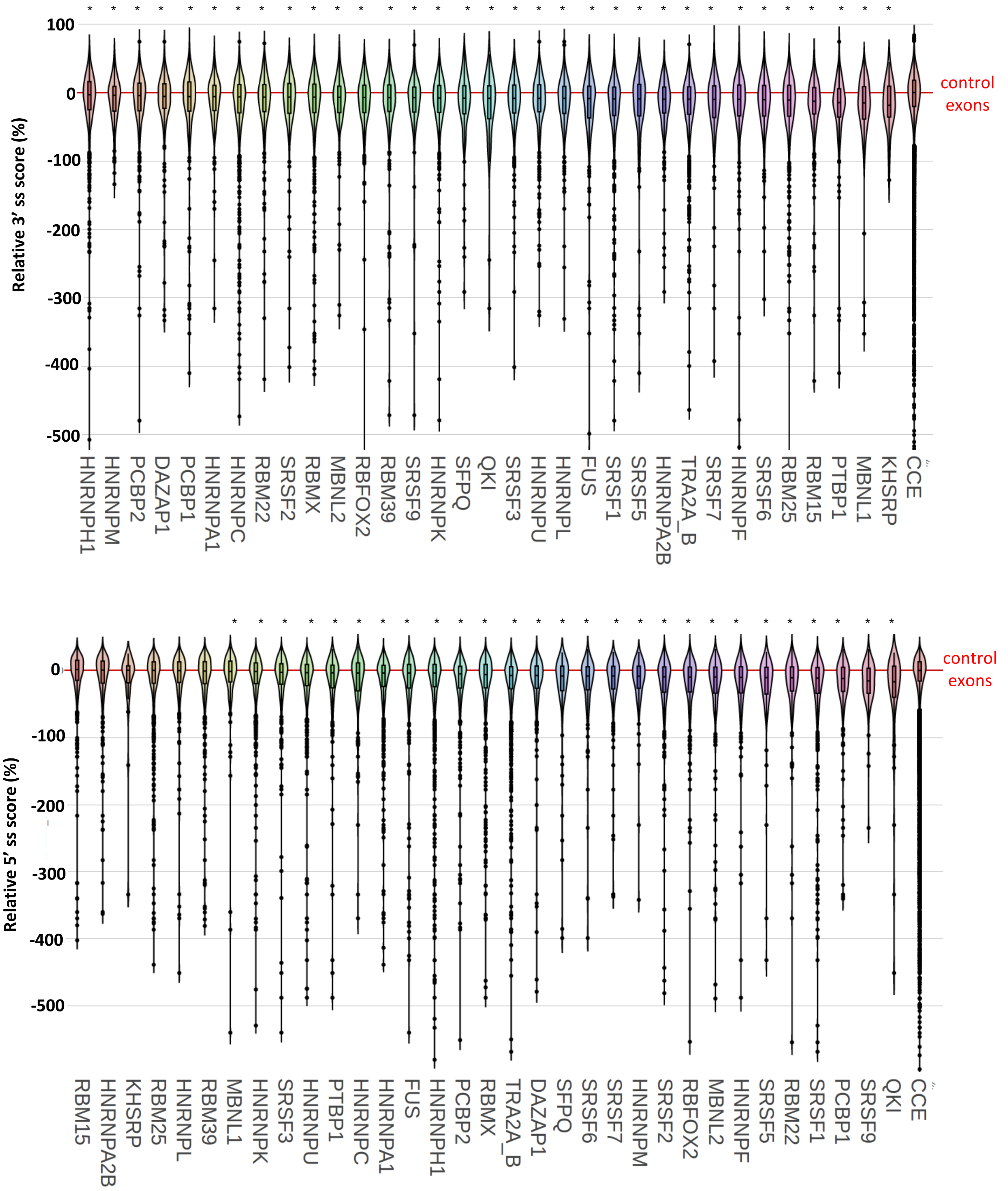
100
0
-100
-200
-300
-400
-500
Relative 3' ss score (%)
control exons
HNRNPH1
HNRNPM
PCBP2
DAZAP1
PCBP1
HNRNPA1
HNRNPC
RBM22
SRSF2
RBMX
MBNL2
RBFOX2
RBM39
SRSF9
HNRNPK
SFPQ
QKI
SRSF3
HNRNPU
HNRNPL
FUS
SRSF1
SRSF5
HNRNPA2B
TRA2A_B
SRSF7
HNRNPF
SRSF6
RBM25
RBM15
PTBP1
MBNL1
KHSRP
CCE
0
-100
-200
-300
-400
-500
Relative 5' ss score (%)
control exons
RBM15
HNRNPA2B
KHSRP
RBM25
HNRNPL
RBM39
MBNL1
HNRNPK
SRSF3
HNRNPU
PTBP1
HNRNPC
HNRNPA1
FUS
HNRNPH1
PCBP2
RBMX
TRA2A_B
DAZAP1
SFPQ
SRSF6
SRSF7
HNRNPM
SRSF2
RBFOX2
MBNL2
HNRNPF
SRSF5
RBM22
SRSF1
PCBP1
SRSF9
QKI
CCE

SUP S2

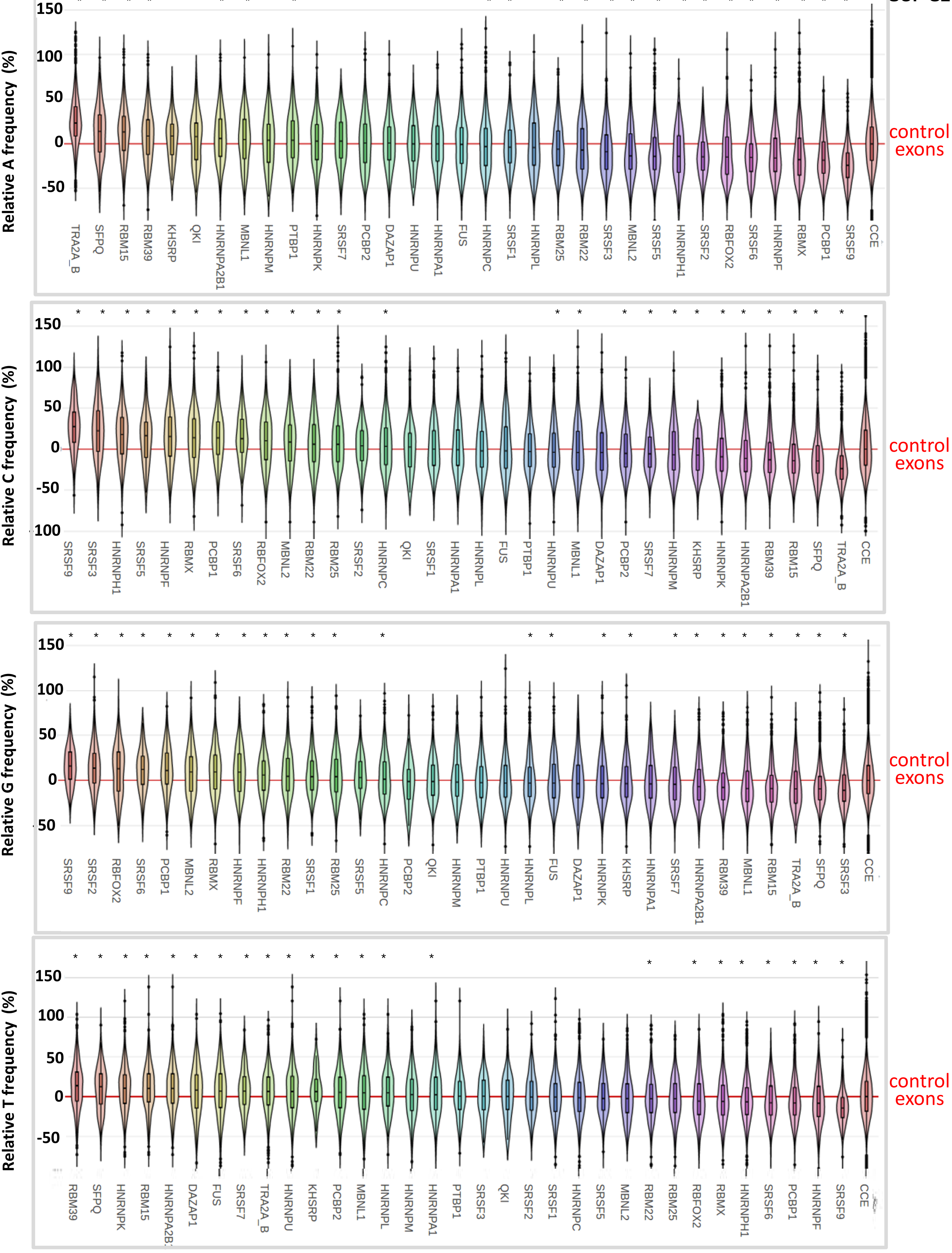
Relative A frequency (%)
150
100
50
0
-50
control exons
TRA2A_B SFPQ RBM15 RBM39 KHSRP QKI HNRNPA2B1 MBNL1 HNRNPM PTBP1 HNRNPK SRSF7 PCBP2 DAZAP1 HNRNPU HNRNPA1 FUS HNRNPC SRSF1 HNRNPL RBM25 RBM22 SRSF3 MBNL2 SRSF5 HNRNPH1 SRSF2 RBFOX2 SRSF6 HNRNPF RBMX PCBP1 SRSF9 CCE
Relative C frequency (%)
150
100
50
0
-50
-100
control exons
SRSF9 SRSF3 HNRNPH1 SRSF5 HNRNPF RBMX PCBP1 SRSF6 RBFOX2 MBNL2 RBM22 RBM25 SRSF2 HNRNPC QKI SRSF1 HNRNPA1 HNRNPL FUS PTBP1 HNRNPU MBNL1 DAZAP1 PCBP2 SRSF7 HNRNPM KHSRP HNRNPK HNRNPA2B1 RBM39 RBM15 SFPQ TRA2A_B CCE
Relative G frequency (%)
150
100
50
0
-50
control exons
SRSF9 SRSF2 RBFOX2 SRSF6 PCBP1 MBNL2 RBMX HNRNPF HNRNPH1 RBM22 SRSF1 RBM25 SRSF5 HNRNPC PCBP2 QKI HNRNPM PTBP1 HNRNPU HNRNPL FUS DAZAP1 HNRNPK KHSRP HNRNPA1 SRSF7 HNRNPA2B1 RBM39 MBNL1 RBM15 TRA2A_B SFPQ SRSF3 CCE
Relative T frequency (%)
150
100
50
0
-50
control exons
RBM39 SFPQ HNRNPK RBM15 HNRNPA2B1 DAZAP1 FUS SRSF7 TRA2A_B HNRNPU KHSRP PCBP2 MBNL1 HNRNPL HNRNPM HNRNPA1 PTBP1 SRSF3 QKI SRSF2 SRSF1 HNRNPC SRSF5 MBNL2 RBM22 RBM25 RBFOX2 RBMX HNRNPH1 SRSF6 PCBP1 HNRNPF SRSF9 CCE

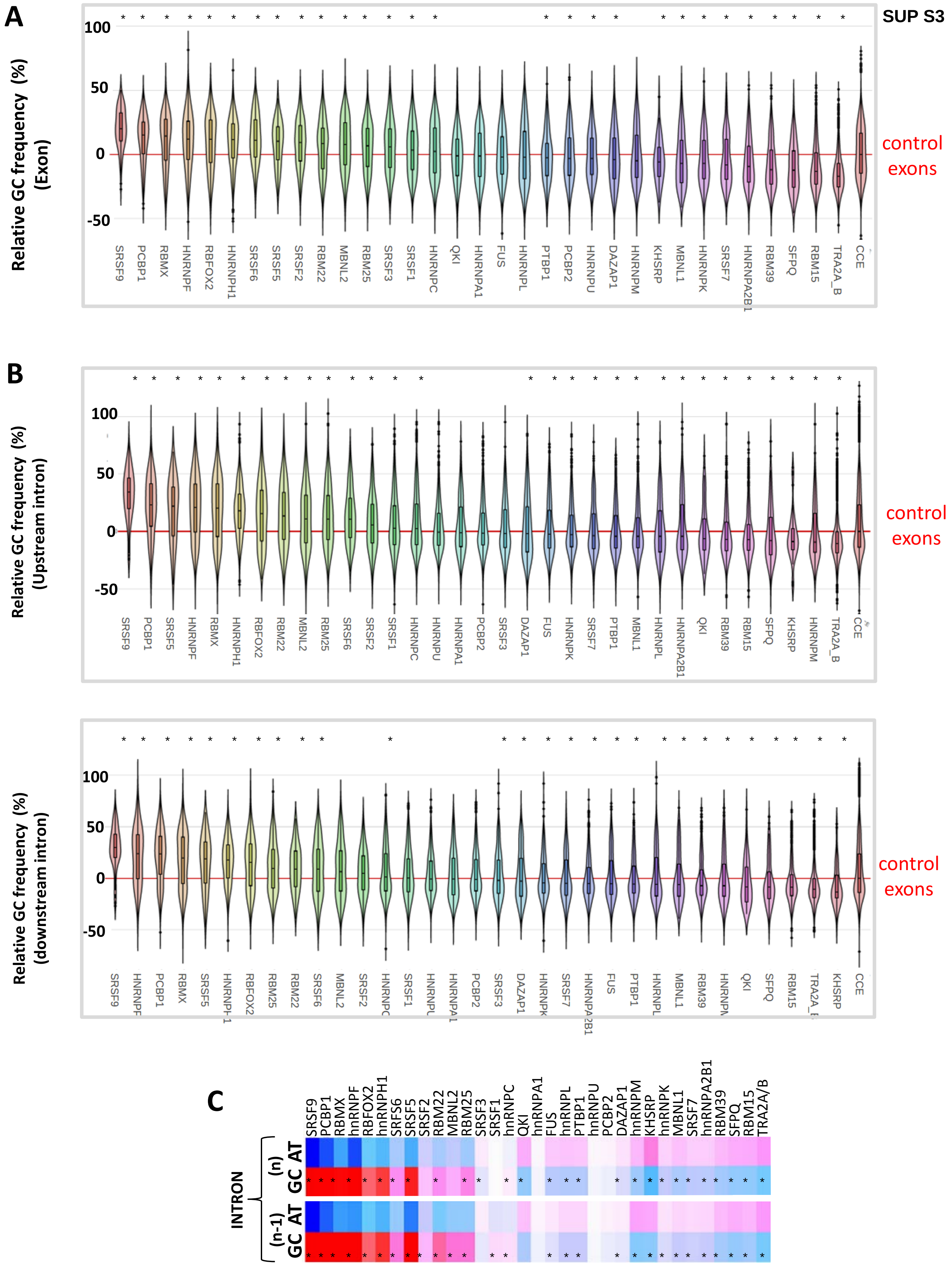

SUP S3
A
Relative GC frequency (%) (Exon)
100
50
0
-50
SRSF9 PCBP1 RBMX HNRNPF RBFOX2 HNRNPH1 SRSF6 SRSF5 SRSF2 RBM22 MBNL2 RBM25 SRSF3 SRSF1 HNRNPC QKI HNRNPA1 FUS HNRNPL PTBP1 PCBP2 HNRNPU DAZAP1 HNRNPM KHSRP MBNL1 HNRNPK SRSF7 HNRNPA2B1 RBM39 SFPQ RBM15 TRA2A_B CCE
control exons
B
Relative GC frequency (%) (Upstream intron)
100
50
0
-50
SRSF9 PCBP1 SRSF5 HNRNPF RBMX HNRNPH1 RBFOX2 RBM22 MBNL2 RBM25 SRSF6 SRSF2 SRSF1 HNRNPC HNRNPU HNRNPA1 PCBP2 SRSF3 DAZAP1 FUS HNRNPK SRSF7 PTBP1 MBNL1 HNRNPL HNRNPA2B1 QKI RBM39 RBM15 SFPQ KHSRP HNRNPM TRA2A_B CCE
control exons
Relative GC frequency (%) (downstream intron)
100
50
0
-50
SRSF9 HNRNPF PCBP1 RBMX SRSF5 HNRNPH1 RBFOX2 RBM25 RBM22 SRSF6 MBNL2 SRSF2 HNRNPC SRSF1 HNRNPU HNRNPA1 PCBP2 SRSF3 DAZAP1 HNRNPK SRSF7 HNRNPA2B1 FUS PTBP1 HNRNPL MBNL1 RBM39 HNRNPM QKI SFPQ RBM15 TRA2A_B KHSRP CCE
control exons
C
SRSF9 PCBP1 RBMX hnRNPF RBFOX2 hnRNPH1 SRFS6 SRSF5 SRSF2 RBM22 MBNL2 RBM25 SRSF3 SRSF1 hnRNPC QKI hnRNPA1 FUS hnRNPL PTBP1 PCBP2 DAZAP1 hnRNPM KHSRP hnRNPK MBNL1 SRSF7 hnRNPA2B1 RBM39 SFPQ RBM15 TRA2A/B
INTRON
(n)
AT
GC
(n-1)
AT
GC

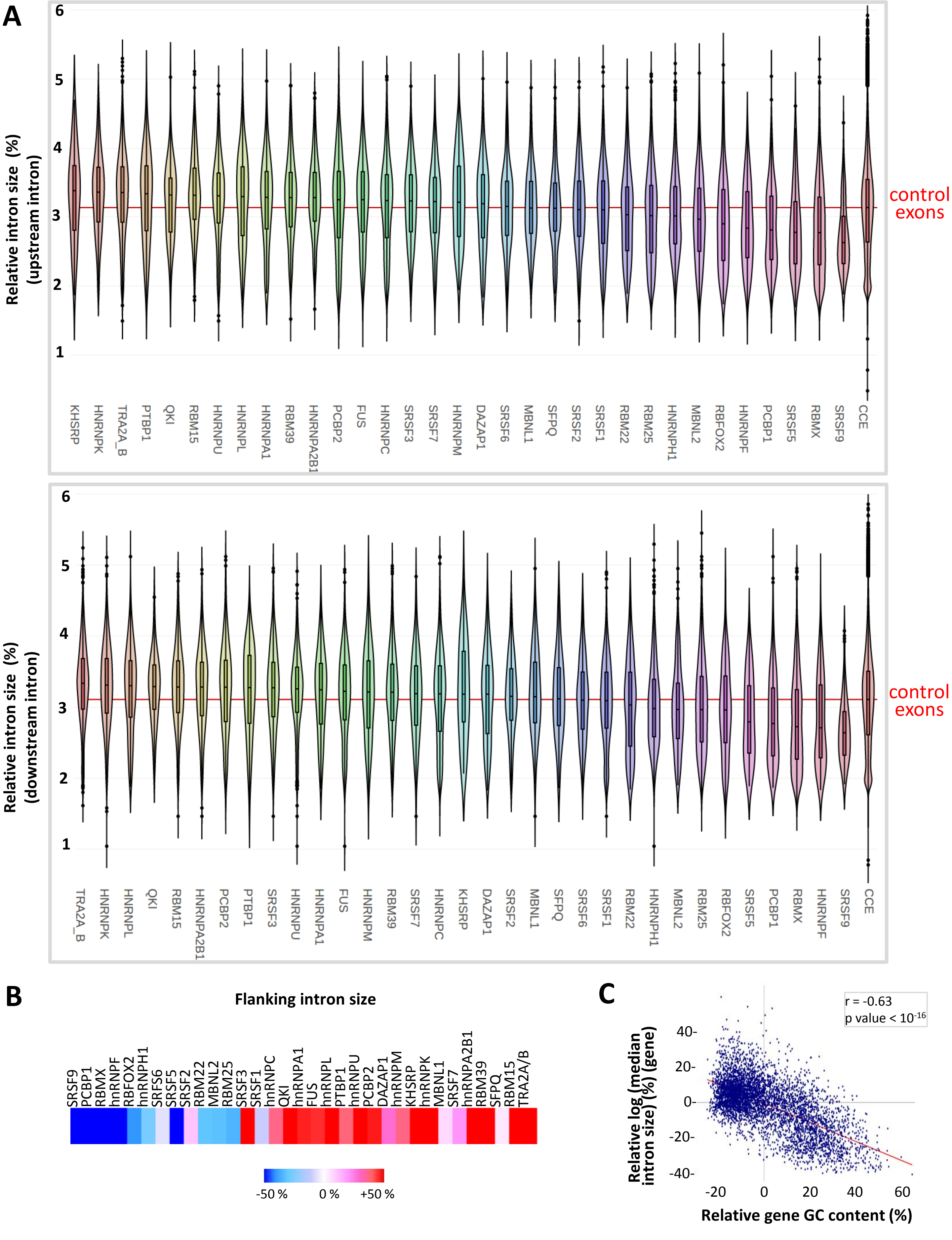
A
Relative intron size (%) (upstream intron)
6
5
4
3
2
1
KHSRP
HNRNPK
TRA2A_B
PTBP1
QKI
RBM15
HNRNPU
HNRNPL
HNRNPA1
RBM39
HNRNPA2B1
PCBP2
FUS
HNRNPC
SRSF3
SRSF7
HNRNPM
DAZAP1
SRSF6
MBNL1
SFPQ
SRSF2
SRSF1
RBM22
RBM25
HNRNPH1
MBNL2
RBFOX2
HNRNPF
PCBP1
SRSF5
RBMX
SRSF9
CCE
control exons
Relative intron size (%) (downstream intron)
TRA2A_B
HNRNPK
HNRNPL
QKI
RBM15
HNRNPA2B1
PCBP2
PTBP1
SRSF3
HNRNPU
HNRNPA1
FUS
HNRNPM
RBM39
SRSF7
HNRNPC
KHSRP
DAZAP1
SRSF2
MBNL1
SFPQ
SRSF6
SRSF1
RBM22
HNRNPH1
MBNL2
RBM25
RBFOX2
SRSF5
PCBP1
RBMX
HNRNPF
SRSF9
CCE
control exons
B
Flanking intron size
SRSF9
PCBP1
RBMX
hnRNPF
RBFOX2
hnRNPH1
SRFS6
SRSF5
SRSF2
RBM22
MBNL2
RBM25
SRSF3
SRSF1
hnRNPC
QKI
hnRNPA1
FUS
hnRNPL
PTBP1
hnRNPU
PCBP2
DAZAP1
hnRNPM
KHSRP
hnRNPK
MBNL1
SRSF7
hnRNPA2B1
RBM39
SFPQ
RBM15
TRA2A/B
-50 %
0 %
+50 %
C
r = -0.63
p value < 10-16
Relative log (median intron size) (%) (gene)
40-
20-
0-
-20-
-40-
-20
0
20
40
60
Relative gene GC content (%)

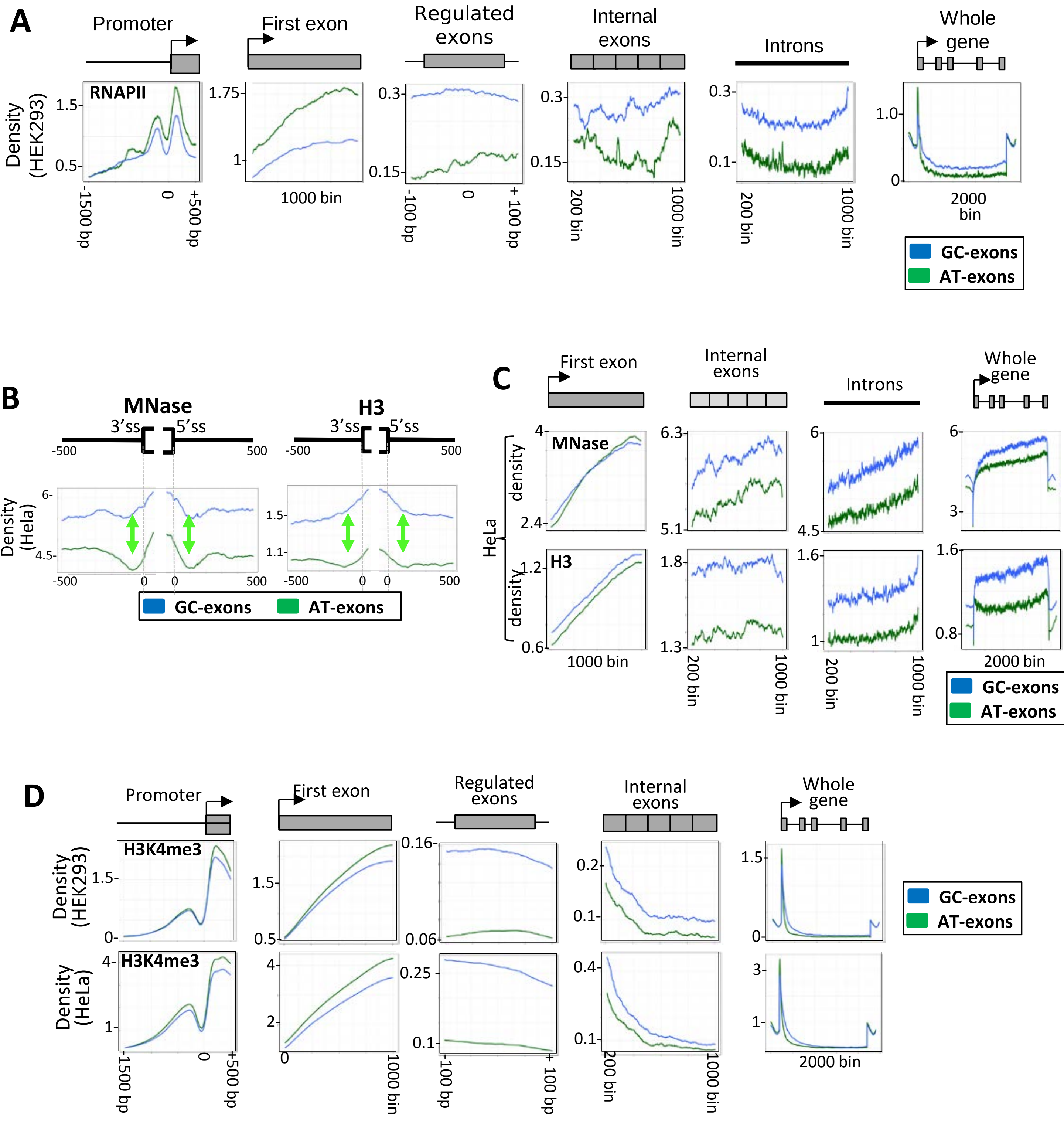
A
Promoter
First exon
Regulated exons
Internal exons
Introns
Whole gene
RNAPII
Density (HEK293)
1.5
0.5
1.75
1
0.3
0.15
0.3
0.15
0.3
0.1
1.0
0
-1500 bp
0
+500 bp
1000 bin
-100 bp
0
+ 100 bp
200 bin
1000 bin
200 bin
1000 bin
2000 bin
GC-exons
AT-exons
B
MNase
3'ss
5'ss
-500
500
H3
Density (Hela)
6
4.5
1.5
1.1
0
C
First exon
Internal exons
Introns
Whole gene
MNase
H3
HeLa
density
4
2.4
6.3
5.1
6
4.5
3
1.2
0.6
1.8
1.3
1.6
1
0.8
D
H3K4me3
Density (HEK293)
Density (HeLa)
0.16
0.06
0.2
0.1
0.25
0.4
2
1.5
0.5
3

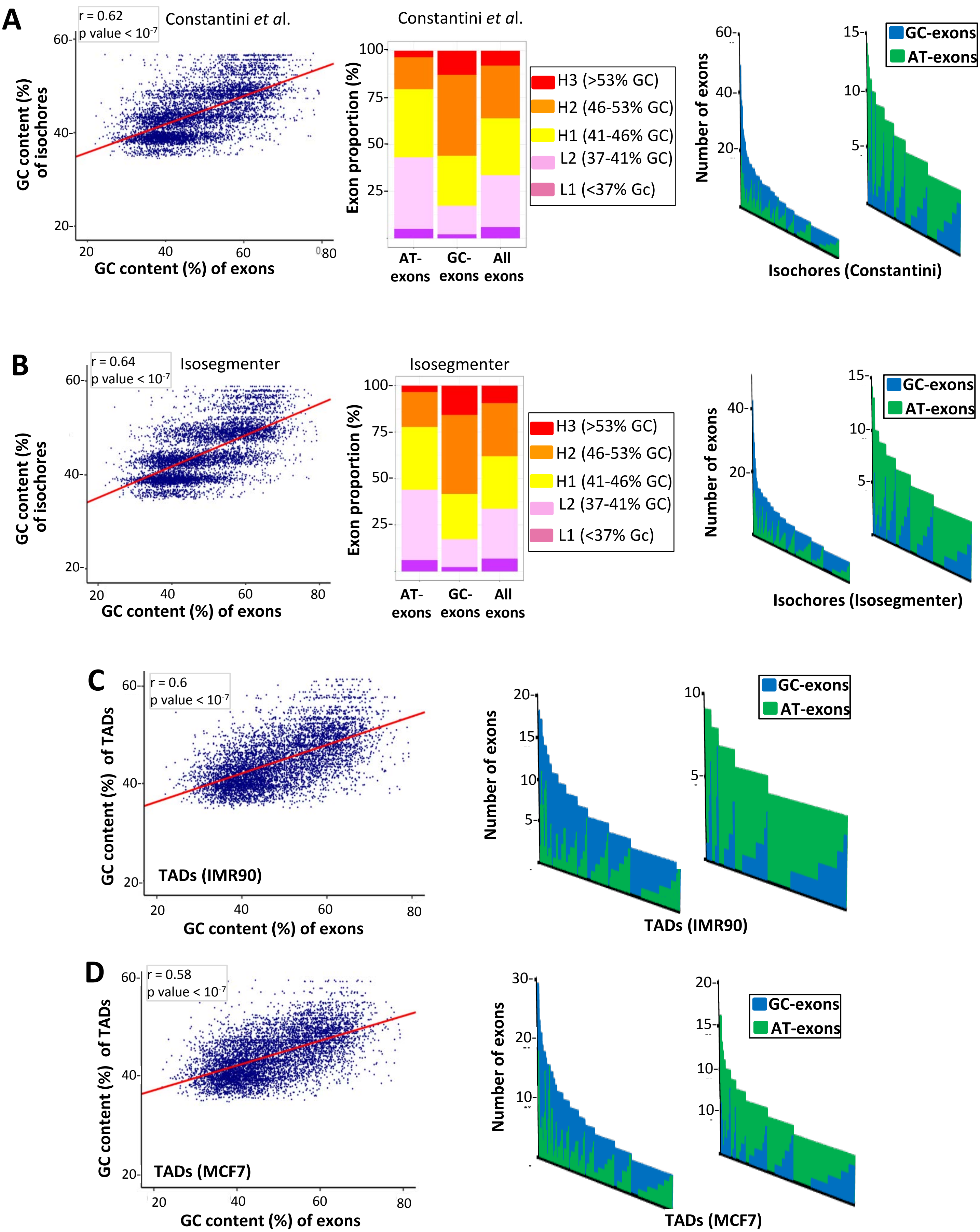
A
r = 0.62
p value < 10-7
Constantini et al.
GC content (%) of isochores
GC content (%) of exons
Constantini et al.
Exon proportion (%)
H3 (>53% GC)
H2 (46-53% GC)
H1 (41-46% GC)
L2 (37-41% GC)
L1 (<37% Gc)
AT-exons
GC-exons
All exons
Number of exons
GC-exons
AT-exons
Isochores (Constantini)
B
r = 0.64
p value < 10-7
Isosegmenter
GC content (%) of isochores
GC content (%) of exons
Isosegmenter
Exon proportion (%)
H3 (>53% GC)
H2 (46-53% GC)
H1 (41-46% GC)
L2 (37-41% GC)
L1 (<37% Gc)
AT-exons
GC-exons
All exons
Number of exons
GC-exons
AT-exons
Isochores (Isosegmenter)
C
r = 0.6
p value < 10-7
GC content (%) of TADs
TADs (IMR90)
GC content (%) of exons
Number of exons
GC-exons
AT-exons
TADs (IMR90)
D
r = 0.58
p value < 10-7
GC content (%) of TADs
TADs (MCF7)
GC content (%) of exons
Number of exons
GC-exons
AT-exons
TADs (MCF7)

**Sup S7**

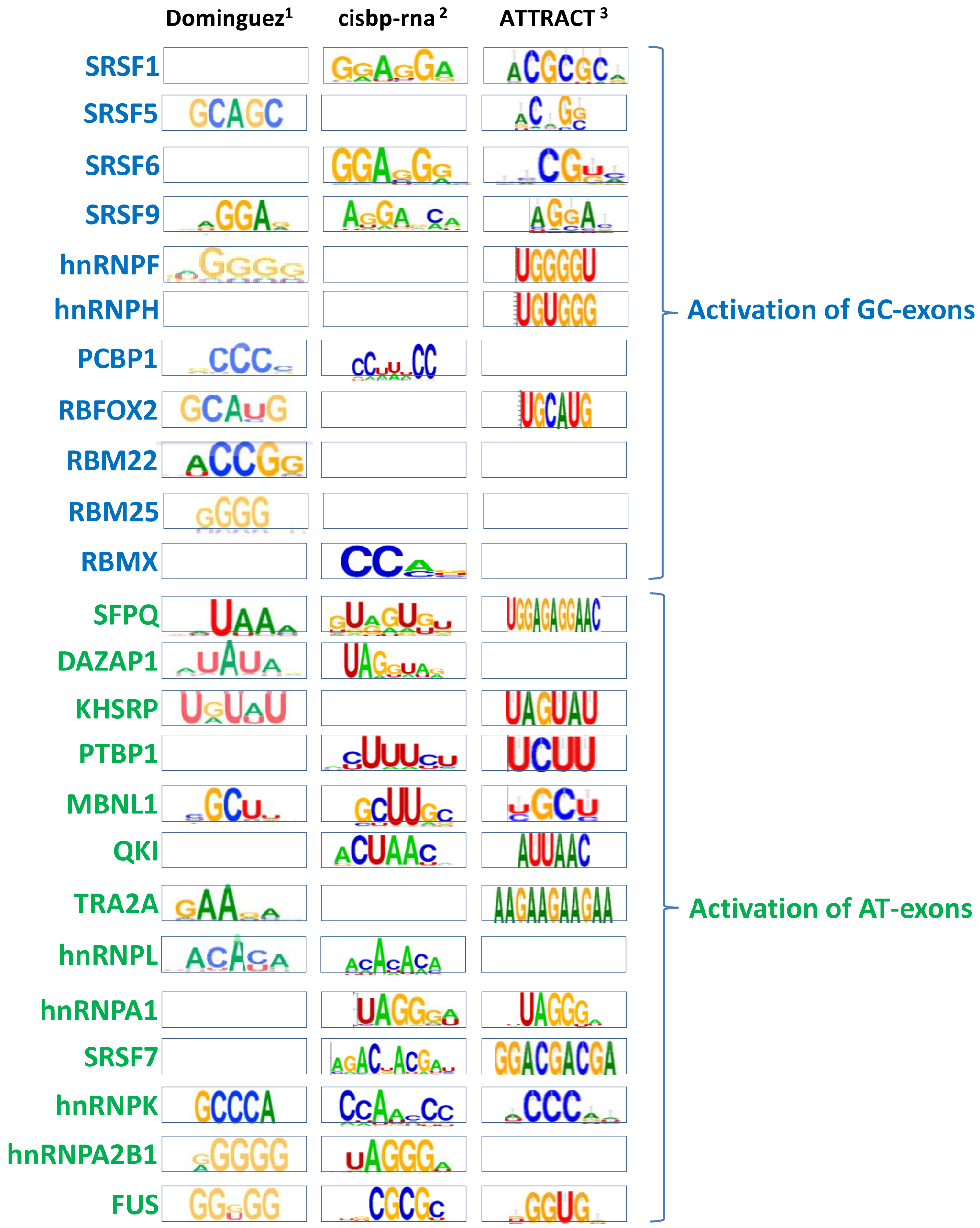


[1] Dominguez et al. PMID 29883606
[2] http://attract.cnic.es
[3] http://cisbp-rna.ccbr.utoronto.ca/index.php
